# Universality of clonal dynamics poses fundamental limits to identify stem cell self-renewal strategies

**DOI:** 10.1101/2020.02.10.941286

**Authors:** Cristina Parigini, Philip Greulich

**Affiliations:** School of Mathematical Science, University of Southampton, Southampton SO17 1BJ, United Kingdom; Institute for Life Sciences, University of Southampton, Southampton SO17 1BJ, United Kingdom

## Abstract

How adult stem cells maintain self-renewing tissues is *in vivo* commonly assessed by analysing clonal data from cell lineage tracing assays. To identify strategies of stem cell self-renewal requires that different models of stem cell fate choice predict sufficiently different clonal statistics. Here we show that models of cell fate choice can, in homeostatic tissues, be categorized by exactly two ‘universality classes’, whereby models of the same class predict, under asymptotic conditions, the same clonal statistics. Those classes relate to generalizations of the canonical asymmetric vs. symmetric stem cell self-renewal strategies and are differentiated by a conservation law. This poses both challenges and opportunities to identify stem cell self-renewal strategies: while under asymptotic conditions, self-renewal models of the same universality class cannot be distinguished by clonal data only, models of different classes can be distinguished by simple means.

## Introduction

Adult stem cells are the key players for maintaining and renewing biological tissue, due to their ability to persistently produce tissue cells through cell division and differentiation (1). For maintaining tissues in a homeostatic state it is crucial that stem cells adopt suited *self-renewal strategies*, a pattern of stem cell fate choices that balances proliferation and differentiation; otherwise, imbalanced proliferation may lead to cancer. Therefore, the understanding and identification of stem cell self-renewal strategies has been a major goal of stem cell biology ever since the discovery of adult stem cells.

Classically, two stem cell self-renewal strategies have been proposed (2, 3): following the *Invariant Asymmetric division* (IA), stem cells undertake only asymmetric divisions, whose outcome is one differentiating cell and one stem cell as daughter cells. The other proposed strategy, *Population Asymmetry* (PA) (2–5), features additionally symmetric divisions, which produce either two stem cells or two differentiated cells as daughters, yet in balanced proportions. Both patterns of cell fate choice leave the number of cells on average unchanged and thus can maintain homeostasis. Assessing stem cell self-renewal strategies experimentally is difficult *in vivo*, since direct observation of cell divisions is rarely possible. Yet, through genetic cell lineage tracing assays, the statistics of clones – the progeny of individual cells – can be obtained, and via mathematical modelling assessing cell fate dynamics became possible. With such an approach several studies suggested that population asymmetry prevails in many mouse tissues (e.g. Refs. (6–10)).

However, the interpretation of those studies has been challenged by a suggested alternative self-renewal strategy, called *Dynamic Heterogeneity* (DH), featuring some degree of cell fate plasticity (11). In this model all stem cell divisions are asymmetric, yet it is in agreement with the experimental clonal data that had previously been shown to agree also with the population asymmetry strategy. Thus, those two strategies are not distinguishable in view of the clonal data.

This raises the question to what extent different stem cell self-renewal strategies can be distinguished at all via clonal data (5, 12). Here, we address this question by studying models for stem cell fate choice, which define the self-renewal strategies, in their most generic form. We show that many cell fate models predict, under asymptotic conditions, the same clonal statistics and thus cannot be distinguished via clonal data from cell lineage tracing experiments. In particular, we find that there exist two particular classes of stem cell self-renewal strategies: one class of models which all generate an Exponential distribution of clone sizes (the number of cells in a clone) after sufficiently large time, and one which generates a Normal distribution under sufficiently fast stem cell proliferation. Crucially, these two classes are not differentiated via the classical definitions of symmetric and asymmetric stem cell divisions, but by whether or not a subset of cells is conserved. These classes thus bear resemblance to “universality classes” known from statistical physics, as suggested in (5). This leads us to a more generic, and in this context more useful, definition of the terms “symmetric” and “asymmetric” divisions. Notably, however, we find that the conditions for the emergence of universality are not always fulfilled in real tissues, which provides chances, but also further challenges, for the identification of stem cell fate choices in homeostatic tissues.

### Strategies for stem cell self-renewal

The two classical stem cell self-renewal strategies, Invariant Asymmetry (IA) and Population Asymmetry (PA) (2–5), are commonly described in terms of two cell types: stem cells (*S*) which can self-renew (i.e. divide without reducing their potential to divide in the future); and differentiating cells (*D*). Both strategies can be expressed in terms of a single parametrized stochastic model, a multi-type branching process (13), defined by the outcomes of cell divisions (the *cell fate choices*)

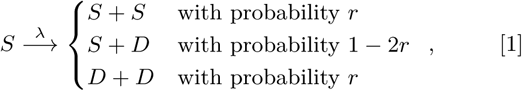

where cells of type *S* divide with rate λ. Here, a daughter cell configuration *S* + *S* corresponds to *symmetric self-renewal division* and *D* + *D* to *symmetric differentiation*, while daughter cells of different type, *S* + *D*, marks an *asymmetric division*. In the basic model version, a cell of type *D* is eventually lost with rate *γ*, 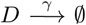 (corresponding to death, shedding, or emigration of *D*-cells), while other versions may include the possibility of limited proliferation as committed progenitor cells. The two self-renewal strategies, IA and PA, are distinguished by the value of the *symmetric division fraction r*: the PA model corresponds to any **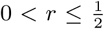** ; the IA model is defined by *r* = 0, i.e. only asymmetric divisions occur. To maintain homeostasis, the number of cells must stay, on average, constant. Thus cells following the PA strategy must regulate the probabilities of symmetric self-renewal and differentiation to be exactly equal, whereas for the IA model this is trivially assured. However, only for the IA model the number of stem cells is *strictly conserved*, i.e. no gain or loss of stem cells is possible.

A way to assess self-renewal strategies experimentally is via genetic cell lineage tracing (14, 15): By marking single cells with an inheritable genetic marker (through a Cre-Lox system (16, 17)) each cell’s progeny, called a *clone*, which retain that marker, can be traced. The number of cells per clone, that is the *clone size*, is measured and the statistical frequency distribution of clone sizes (*clone size distribution*) determined. To test the cell fate choice models on that data, one evaluates the models with a single cell as initial condition and samples the outcome in terms of the final cell numbers – the size of a virtual clone. In the basic version of the model (i.e. when 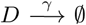), the IA and PA models predict, respectively, a Poisson and an Exponential clone size distribution for large times (5, 18) (see also the SI, section 3.A). Thus, they are fundamentally different and can easily be distinguished when compared with clonal data. By a series of lineage tracing experiments it was confirmed that Exponential clone size distributions prevail for most mouse tissues, which thus exclude the IA model and support the PA strategy (6–10).

While this seemed to settle the case in favour of the PA strategy, at least for most adult mouse tissues, this was challenged by a third type of strategy, the DH model (11). Motivated by the emerging view of prevailing cell plasticity (19–22), the DH model considers the possibility of reversible switching between two cell types:

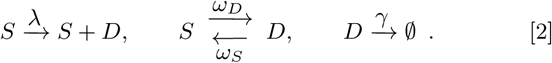

where symbols at arrows denote the *process rates* (frequency of events). This strategy is also capable to maintain a homeostatic population if *γ/*λ = *ω*_*S*_ */ω*_*D*_. Notably, the DH model only features asymmetric divisions (in that daughter cells are of different type), like the IA model, yet the DH model predicts clonal statistics that are indistinguishable from the PA model (11). This means that in view of the existing clonal data for mouse tissues, the DH model, may as well describe the real cell fate dynamics. More fundamentally, this implies that the PA and DH model cannot be distinguished via plain clonal data, which poses fundamental limitations to the common approach to use lineage tracing for determining cell fate choices.

This demonstrates that the classical definition of asymmetric and symmetric divisions is not always suitable to distinguish cell fate strategies in view of clonal data alone. In general, cell fate dynamics may be much more complex than the simplified models described above, as there may exists a plethora of cell (sub-)types in a tissue. However, to what extent would it be possible to distinguish details of potentially rather complex cell fate dynamics models through comparison with clonal data at all? This is only the case if the clonal statistics are sufficiently different. In the following we study cell fate models in their most generic form, and analyze what clonal statistics would be expected.

## Results

### Model Generalization

Let us consider the dynamics of a generic system of cells, characterized by a number *m* of possible cell states *X*_*i*_, *i* = 1, …, *m*. We define a cell *state* here as a group of cells showing common properties (e.g. any cell sub-type classification). Most generally, cells in a state *X*_*i*_ may be able to divide, producing daughter cells of any cell states *X*_*j*_ and *X*_*k*_ (where *i* = *j* = *k*, i.e. simple cell duplication, is possible). Furthermore, any cell state *X*_*i*_ may turn into another state *X*_*j*_ or may be lost (through emigration, shedding, or death). Hence, we can write a generic cell fate model as

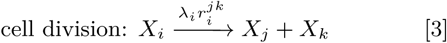

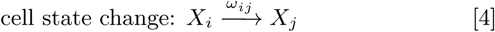

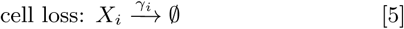

where *i, j, k* = 1, …, *m*. In this model, λ_*i*_ is the rate of division of cells in state *X*_*i*_ and the parameter 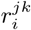 corresponds to the proportion of division outcomes producing daughter cells of state *X*_*j*_ and state *X*_*k*_; *ω*_*ij*_ is the transition rate from state *X*_*i*_ to state *X*_*j*_ and *γ*_*i*_ the loss rate from state *X*_*i*_. This model is a general multi-type branching process, which is suitable to describe cell population dynamics in any dimension larger than one, even under cell-extrinsic regulation (5, 23).

In the following, we study the dynamics of cell numbers in each state *X*_*i*_, *n*_*i*_. To gain initial insight into those dynamics, let us first consider the time evolution of the *mean* cell numbers, 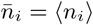, given by,

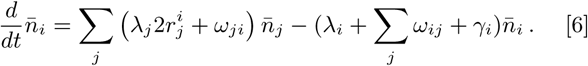

in which 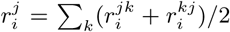 is the probability of having a daughter cell in state *X*_*j*_ produced upon division of a cell in state *X*_*i*_. This linear system of differential equations can be written more compactly in terms of the mean cell number vector 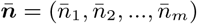,

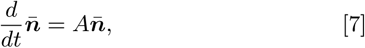

with *A* being the *m* × *m* matrix

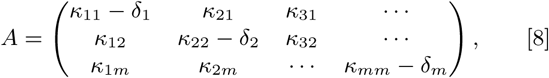

where we defined the *total transition rate* 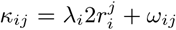, combining all transitions from *X*_*i*_ to *X*_*j*_ by cell divisions and direct transitions, and the *local loss rate δ*_*i*_ = λ_*i*_ + ∑_*j*_ *ω*_*ij*_ + *γ*_*i*_.

Models of the form 3-5 are not generally in homeostasis, which in this context is defined by the existence of a stationary state 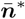, with 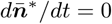, that is stable and non-trivial^∗^. This can in principle be assessed through the spectral properties of *A* (24), but applying spectral conditions explicitly is unwieldy and difficult to interpret biologically. For a more intuitive view, we interpret the system, Eq. 7, as a network (graph): the matrix *A* can be interpreted as the adjacency matrix of the *cell state network*. This is a weighted directed graph in which cell states correspond to the graph’s nodes and a link from state *X*_*i*_ to *X*_*j*_ exists where a transition is possible, i.e. when *κ*_*ij*_ > 0. The value of *κ*_*ij*_ also denotes the link weights (diagonal elements of *A* can be considered as self-links). Now, we note that Eq. 7 is linear and cooperative, i.e. the off-diagonal elements of matrix *A* are non-negative, and for such systems more simple and intuitive conditions for homeostasis exist (25), based on a decomposition into the network’s *Strongly Connected Component (SCC)*. An SCC is a sub-graph that groups nodes which are *strongly connected*, i.e. which are mutually connected by paths (more accurately: two nodes, *X*_*i*_ and *X*_*j*_ are strongly connected if there exists a path from *X*_*i*_ to *X*_*j*_ and from *X*_*j*_ to *X*_*i*_ on the network). An example of such a decomposition, which yields an *acyclic* condensed network that contains SCCs as nodes and directed links between them, is shown in Fig. 1.

**Fig. 1.**
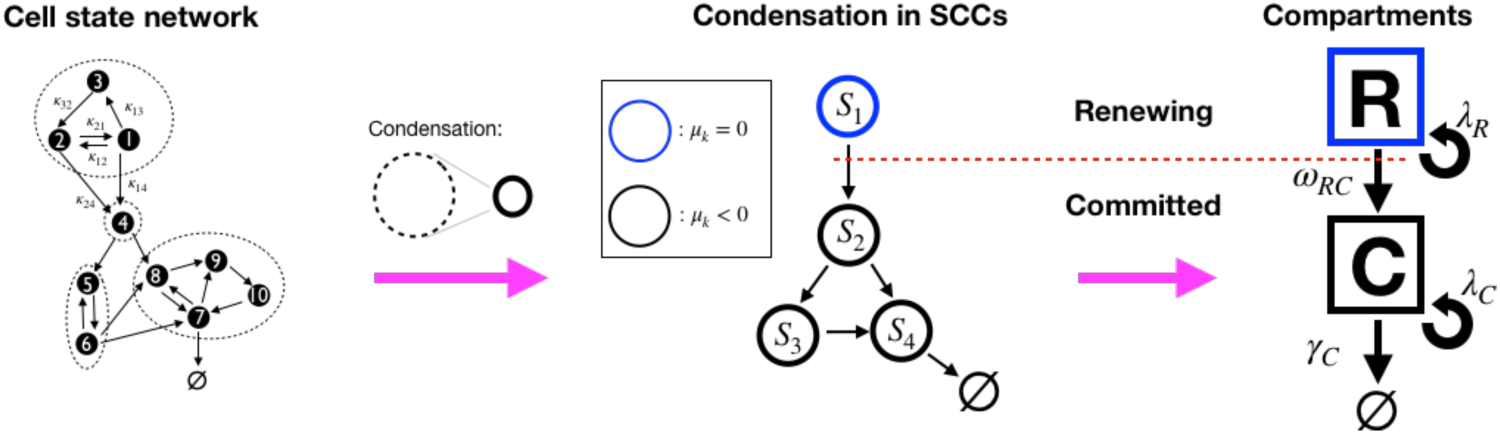
Illustration of the decomposition of a homeostatic cell state network into SCCs and the compartment representation, Eq. 9. (Left): An example cell state network representing the matrix *A* in Eq. 8 (self-links not displayed). The dashed circles denote the network’s Strongly Connected Components (SCCs) (see definition in text). (Middle): The *Condensed network* is the corresponding network of SCCs, *S*_*k*_, wherein SCCs are the nodes and a link between two SCCs exists if any of their states are connected. For homeostatic networks, an SCC with dominant eigenvalue *µ* = 0 is at the apex, while other SCCs have *µ* < 0. (Right): We distinguish two compartments, the Renewing compartment ℛ, consisting of the apex SCC, with *µ* = 0, and the Committed compartment *C* consisting of the remainder, with *µ* < 0.

The stability of systems like Eq. 7 is then determined by the dominant eigenvalues *µ*_*k*_ of each strongly connected component *k*, for *k* = 1, …, *m*_*S*_ where *m*_*S*_ is the number of SCCs^†^, and their topological arrangement. In brief, according to Ref. (25), the conditions for existence of a homeostatic state are that, at the apex of each lineage (the condensed cell state network), there must be an SCC with dominant eigenvalue *µ*_*k*_ = 0, while all SCCs downstream of the former must have *µ*_*k*_ < 0 (see detailed discussion in the SI, section 1). Given this structure of homeostatic models, we can define two compartments in the cell state transition network: (1) the (self-)**R**enewing compartment (ℛ), which is the SCC at the apex of the lineage tree; and (2) the **C**ommitted compartment (𝒞), which consists of all SCCs with *µ*_*k*_ < 0, i.e, those downstream of the apex SCC. Importantly, cells in states forming ℛ have the potential to return to any state within the same compartment and this population maintains itself. Instead, the cell population in 𝒞 would vanish without external input, since the combined dominant eigenvalue of all those SCCs is negative (it is the union of all SCCs’ *µ*_*k*_ < 0), thus the progeny of each cell in the committed compartment will eventually be lost. We can thereby classify cells as being of a (self-)*Renewing type* (*R*) if their state is within ℛ, and of a *Committed type* (*C*) if their state is in 𝒞. With this coarse-grained classification, a generic homeostatic model can be represented in terms of compartments ℛ and 𝒞 as,

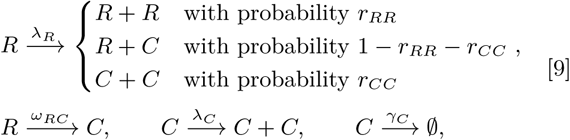

where the symbols above arrows are the *effective rates* of those events, denoting the average frequency at which they occur (loss events *R* → ∅ are not explicitly included, since they can be approximated by a short lived state *X*_*d*_ in 𝒞, as *R* → *X*_*d*_ → ∅). To be compatible with a homeostatic condition, it is further required that (i) the *R*-population remains on average constant (*µ*_*k*_ = 0), i.e. λ_*R*_*r*_*RR*_ = λ_*R*_*r*_*CC*_ + *ω*, and (ii) the loss rate of *C* must exceed its proliferation rate (*µ*_*k*_ < 0), i.e. *γ*_*C*_ > λ_*C*_. Figure 1 shows how a generic homeostatic cell state network can be condensed into an effective model of renewing and committed cell states, according to Eq. 9. It has to be noted, however, that the events depicted in Eq. 9 are *not Markovian*, i.e. the timing of events is not independent from each other and depends on their history. Thus, the ‘rates’ λ_*R*_, λ_*C*_, *ω*_*RC*_, and *γ*_*C*_ are not constant rates in the Markovian sense, yet we can define them by the mean frequency of events occurring.

The formulation in terms of renewing and committed states can help us to gain insights into potential behaviours of generic homeostatic cell fate models. In particular, we define *generalized asymmetric divisions* as events of the type *R → R* + *C*, and *generalized symmetric divisions* as events of the type *R* → *R* + *R* (symmetric renewal) and *R* → *C* + *C* (symmetric commitment). With these definitions, we can categorize homeostatic cell fate models into two classes: *Generalized Invariant Asymmetry* (GIA) models are those which only exhibit *R* → *R* +*C* divisions in the renewing compartment, while *Generalized Population Asymmetry* (GPA) are models for which such restriction does not hold. We note that the two classes are equivalently characterized by a conservation law: For GIA models, the number of cells in ℛ is strictly conserved, while for GPA models, such conservation law do not hold^‡^. Naturally, the previously discussed IA model is a GIA model and the PA model is a GPA model. Notably, the DH model (Eq. 2) is of the GPA category, since in that model *S* and *D* cells form a single SCC at the apex of the lineage hierarchy, and thus they are both part of ℛ. Therefore, a division *S* → *S* + *D* in the DH model, which is asymmetric in the conventional sense, corresponds to *R* → *R* + *R* in terms of compartments (Eq. 9) and thus it is a generalized symmetric division. According to this classification, PA and DH modes are both in the same category (GPA), and indeed, both predict the same type of clone size distribution, an Exponential one (11).

### Numerical simulation of random cell fate models

To check whether the correspondence between model class, GIA vs. GPA, and predicted clonal statistics holds in general, we analyze the clonal dynamics numerically, by generating and testing a large number of random stochastic models, implemented via random generation of the parameters λ_*i*_, *ω*_*ij*_, *γ*_*i*_ and *r*^*jk*^. To simulate clones, we perform stochastic simulations based on the Gillespie algorithm (27), following the rules of Eq. 3-5. We run, for each model, a large number of simulations with initially one cell in the compartment, thus the cell population of each simulation run represents one clone. Then we sample their outcomes, the total cell numbers per clone (the *clone size*) *n* = ∑ _*j*_ *n*_*i*_, to obtain predictions for clonal statistics, namely the frequency distribution of clone sizes (*clone size distribution*) and mean clone sizes (see Materials and Methods).

We first study the mean clone size of surviving clones (with *n* > 0), 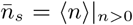, shown in Figure 2, respectively, for the GIA and GPA models, as function of time (the final time *τ* = 20*/α*_min_ where *α*_min_ is the minimal process rate, *α*_min_ = min(λ_1_, …, *ω*_12_, …, *d*_*m*_)). We note that indeed a common behaviour is seen in each case. While for every simulated GIA model, 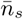 saturates at a plateau value, it steadily increases for every GPA model. This is expected, and can be understood given that clones in a GPA model can go extinct while those in a GIA model not. Assume that there are initially a large number *N*_*c*_ of clones, such that the total number of cells is 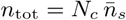. Since the system is homeostatic, it will reach a constant steady state 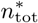 after a sufficient amount of time, meaning that the mean clone size is 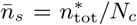. If no clones go extinct, as in GIA models, *N*_*c*_ is constant and thus 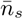 approaches a constant. However, in non-conserved multi-type branching processes, as GPA models are, the clone number *N*_*c*_ decreases through progressive extinction of clones (13), and therefore 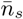 increases, despite the cell population as a whole staying stationary.

**Fig. 2.**
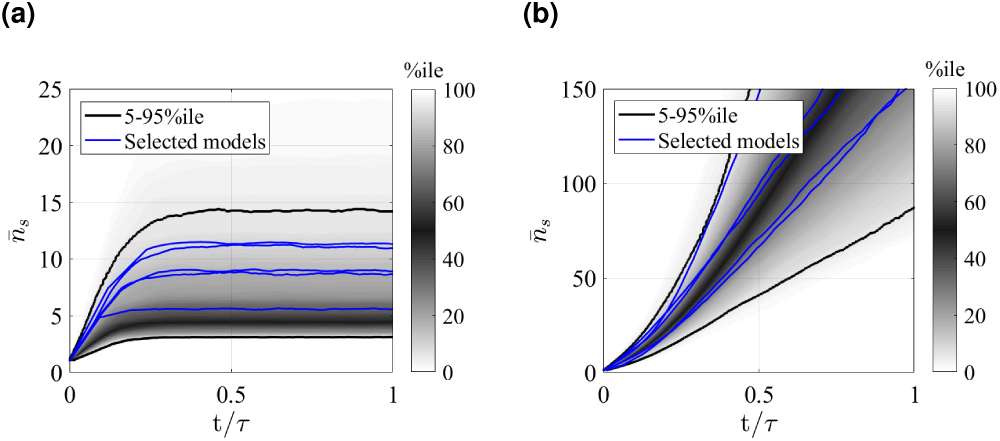
Mean size of surviving clones, 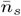, as function of time for random GIA models (a), and GPA models (b). In (a), *τ* = 20*/α*_min_, in (b), *τ* ia the time at 98% clone extinction. The grey shade represents the percentile of all the simulations (black lines limit the 5-95%ile range); the blue curves correspond to some illustrative selected simulations. Simulations for which the final mean is below 2 and where the final condition is not achieved (due to computational limitations) are not included: this results in 238 and 571 models, respectively for the GIA and GPA cases.

The resulting clone size distributions for the two model classes are shown in Figure 3. Here, clones sizes *n* are rescaled by the mean value 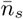 and compared to an Exponential distribution of unitary mean (red curve). As conjectured, all simulated GPA models shown in panel (b) predict asymptotically the same rescaled clone size distribution, namely a standard Exponential distribution. Deviations exist for small times and small clone sizes, but these deviations vanish in the large time limit (details on the convergence are shown in the SI, section 5). This means that different models within the GPA class cannot be distinguished in the long term limit, since they differ only by the mean clone size, which is a free fit parameter. In analogy to statistical physics, we can categorize them as a *universality class* (5), meaning that the details of the model do not affect the (scaled) outcomes for asymptotic conditions, which is a form of weak convergence of random variables (28). However, the same cannot be said about the GIA models. In fact, we see all kind of shapes in the clone size distributions, both peaked distributions and non-peaked ones, and in fact, some distributions are even close to an Exponential form, and can thus not be distinguished from GPA models. The question is whether we can yet find other parameters for which, when large, also GIA models exhibit universality, i.e. yield the same rescaled clone size distribution. For this purpose, we will in the following sections develop a deeper theoretical understanding of the model classes.

**Fig. 3.**
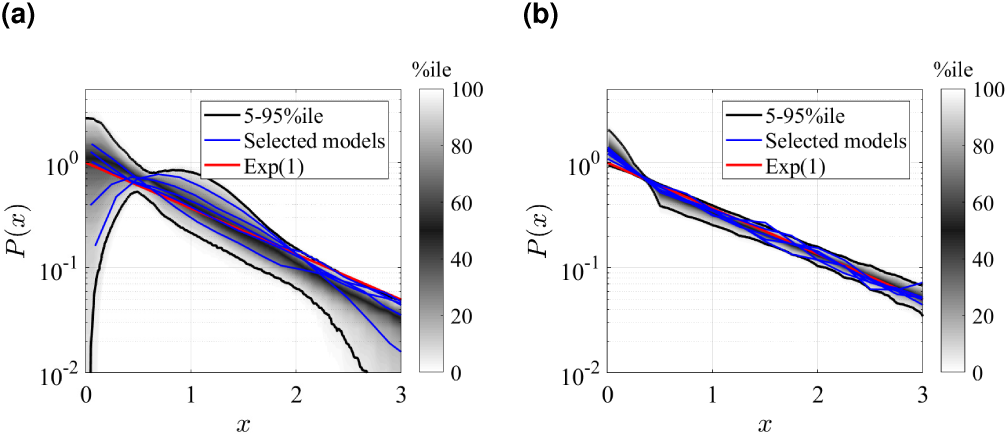
Rescaled clone size distributions (expected relative frequency *P* of clone sizes) for random GIA models (a), and GPA models (b), in terms of the rescaled clone size 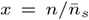, at final time *t* = *τ* (see Fig. 2 for definition). The grey shade represents the percentile of all the simulations (black lines limit the 5-95%ile range); the blue curves correspond to some selected simulations. A reference curve corresponding to an Exponential distribution of unitary mean (‘Exp(1)’) is shown in red.

### Mathematical analysis: Markovian approximation of compartment model

To obtain a deeper understanding of the numerical results, we study the cell fate models in terms of the compartment representation, Eq. 9. In this representation models are not Markovian, yet we can study their Markovian counterpart, as an approximation. While this is not expected to yield accurate clone size distributions in general, the limiting distributions of non-Markovian processes are commonly well estimated by their Markovian counterparts.

For GIA models, which only feature *R* → *R* + *C* transitions between the renewing compartment, ℛ, and the committed compartment, 𝒞, a corresponding Markovian model reads,

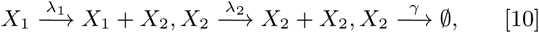

in which *X*_1_ represents a single state in ℛ and *X*_2_ in 𝒞, and symbols at arrows are the process rates. The number of cells in *X*_1_, *n*_1_, is conserved, i.e., given an single *X*_1_-cell initially, it always remains at *n*_1_ = 1. Thus, we only need to consider the dynamics of cells in *X*_2_, *n*_2_. This Markov process can be solved analytically, and for sufficiently large steady state mean number of *X*_2_-cells, 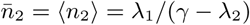 (see SI, section 4.A), the rescaled distribution of cells in *X*_2_ is,

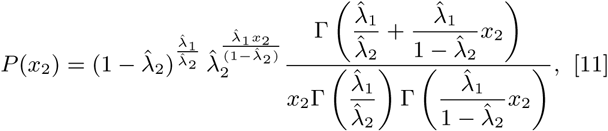

in which 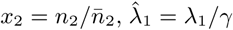 and 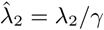. We note that this distribution exhibits a large variety of shapes: for large 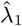 the distribution is peaked, while for small 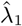 is loses its peak. Notably, for 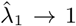 and 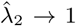, the distribution becomes Exponential and in this case it cannot be distinguished from the GPA case. On the other hand, for 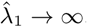, i.e. when the ratio of asymmetric divisions over the loss rate is high, this distribution tends to a Normal distribution with unitary mean and variance equal to 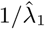. These different behaviours are graphically shown in the SI (see Figures S6, S7 and S8).

For the GPA models, a Markovian approximation reads, accordingly,

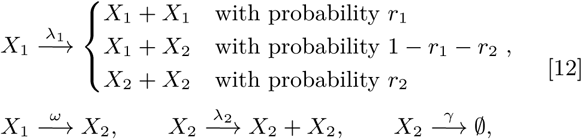

whereby for homeostasis to prevail, λ_1_*r*_1_ = λ_1_*r*_2_ + *ω* and λ_2_ < *γ* must hold. We note that the dynamics of *X*_1_ are independent of *X*_2_ and thus for the number of cells in *X*_1_ in homeostasis holds

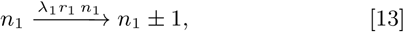

which corresponds to a simple continuous-time branching process with two offspring, for which it is known that the resulting distribution of cell numbers is Exponential, i.e. 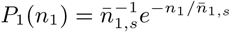, where 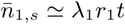 is the mean number of cells in the surviving clones (13).

*X*_2_ cells produced according to 12 follow the same fate as in the 2-state GIA model above. While it is not assured that the distribution of *X*_2_ cells is identical to that of Eq. 11 (due to simultaneous production events of type *X*_1_ → *X*_2_ + *X*_2_), we show in the SI, section 6, that for large rates of production of *C*-cells, the distribution of *C*-cells – here: cells in state *X*_2_ – attains a Normal distribution with mean 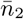 equal to its variance 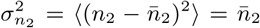. As each *X*_1_ cell contributes independently to the production of *X*_2_-cells, we have that 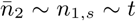. Crucially, this means that in terms of the rescaled variable 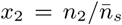 the standard deviation 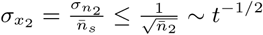 vanishes for large times, since 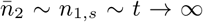. Hence, given fixed *x*_1_, *x*_2_ can be approximated by a constant random number 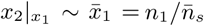. Therefore, the rescaled distribution of the total number of cells is *P* (*x*) = *P*_1_(*x − x*_2_) = *e*^−*x*^, where 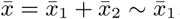. Thus, the rescaled distribution of the total clone size, 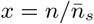, is as well an Exponential.

### Universality of generic cell fate models

For generic GIA or GPA models, the compartment representation, Eq. 9, is not Markovian and one would not expect exactly the distributions we found in the previous section. Fortunately, the limiting distributions of non-Markovian processes and their Markovian counterparts are often, under certain conditions on the parameters, the same. While we reserve the technical arguments for the SI (section 6), we note that this independence of the limiting distribution on the Markov property related to the central limit theorem, which does not rely on the Markov property.

To identify the correct limiting parameters for more complex cell fate models, we need to express the effective non-Markovian rates (i.e. the mean frequency of events) of representation 9 in terms of the original model, 3-5. As discussed in the SI (sections 4.B and 6), we identify those effective rates by the total rates of cell divisions, 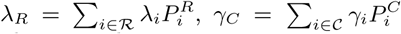, and 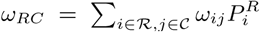 where, for each compartment, 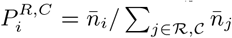 is the probability of a single cell being in state *X*_*i*_ of ℛ, 𝒞, respectively (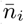 are the solutions to Eq. 6). In the SI, section 6, we reason that all GPA models are expected to generate Exponential clone size distributions for large times *t*. This is indeed what is observed in Fig. 3(b). Correspondingly, for GIA models we expect that for large 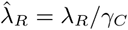 the clone size distribution of GIA models would tend to a Normal distribution. To test this prediction we simulated the same GIA models as for Fig. 3 before, but we tuned parameters in ℛ such that the effective parameter 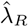 becomes large (see details in the SI, section 4.C). The result is shown in Fig. 4: for an illustrative case shown in panel (a), increasing 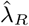 changes the distribution from an exponential form to a peaked form akin to a Normal distribution, and for all simulated random GIA models, shown in panel (b), a Normal distribution is approached when 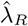 becomes large.

**Fig. 4.**
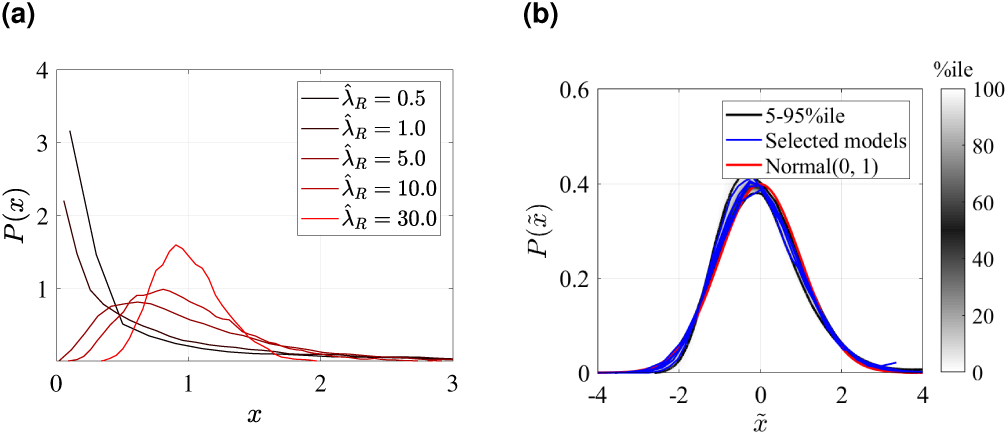
Rescaled clone size distributions (expected relative frequency *P* of clone sizes) for random GIA models as in Fig. 3, at time *t* = *τ* (see definition in Fig. 2). Sensitivity to parameter 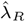 is shown for one illustrative case in panel (a), and all GIA models for 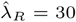 in panel (b). The distributions are shown in terms of the rescaled variables 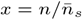 for panel (a) and 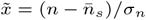, where *σ*_*n*_ is the distributions variance, in panel (b). In (b), the grey shade represents the percentile of all simulations (black lines limit the 5-95%ile range); the blue curves correspond to some selected simulations. A reference curve corresponding to a Normal distribution of zero mean and unitary variance is shown in red. Simulations for which *t* = *τ* is not reached (due to computational limitations) are not included, resulting in 922 model instances.

We note that when taking the limit of large 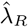, as shown in Fig. 4, also all other process rates *ω*_*ij*_ with *i, j* within ℛ increased as well. What if instead some process rates in ℛ do not scale to become large with 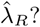 To assess this situation we studied a simple test case similar to model 10 but containing two states in ℛ, connected via direct state transition (see SI, section 4.D). As discussed there, if all rates within ℛ are large compared to the rates in 𝒞 then indeed we observe a Normal clone size distribution, as expected. However, if the direct transition rates between the states of ℛ are smaller or of equal magnitude as *γ*_*C*_, and in addition, one of the two division rates is higher then the other, then we observe a bimodal clone size distribution. The reason is that if the transitions between the two states in ℛ are rare compared to the life time of cells, 1*/γ*_*C*_, they become essentially separated and each of those states generate separate Normal distributions with different mean (due to different cell division rates in those two states) which, when overlaid, generate a bimodal clone size distribution (see detailed arguments in the SI, section 6).

Finally, from those considerations follows:

1. GPA models attain an Exponential clone size distribution for time *t* → ∞.
2. GIA models attain a Normal clone size distribution if all process rates within ℛ are much larger than the inverse lifetime of *C*-cells, *γ*_*C*_.

Hence, the GIA and GPA model classes, each represent a universality class, i.e. a scaling limit exists in which all models of the same class yield the same rescaled clonal statistics.

## Discussion

Our analysis shows that intrinsic limitations exist for identifying strategies of stem cell self-renewal through clonal data from cell lineage tracing experiments. This is due to different models of cell fate choice generating the same type of clonal statistics (clone size distributions), so that model inference based on clonal statistics – currently still the most prevalent method to determine stem cell self-renewal strategies – fails to distinguish them. The feature that different models asymptotically generate the same statistics is a form of weak convergence of random variables (28) and corresponds to *universality*, as known from statistical physics.

Cell fate models can in principle be very complex, with a plethora of cell (sub-)types in a tissue. We introduced a new categorization of cell types, distinguishing between cell states that are committed (*C*-cells), whose progeny is inevitably lost eventually, and non-committed or (self-)renewing cell states (*R*-cells), which retain the potential to remain or return to the apex of the lineage hierarchy. According to this categorization we classified generic models of cell fate choice (for higher than one dimension^§^) as *Generalized Invariant Asymmetry* (GIA), if only generalized asymmetric divisions of the form *R* → *R* + *C* occur for *R*-cells, and *Generalized Population Asymmetry* (GPA), when all kind of divisions can occur, as long as gain and loss of *R*-cells are balanced. Models of the GIA category are also characterized by a conservation law, since the number of *R*-cells is strictly conserved, while GPA models do not exhibit such a conservation law. We found that this classification of models mirrors the clonal statistics generated by them: models of the GPA class all generate clonal statistics which with time converge to an Exponential clone size distribution. Thus, two GPA models can therefore not be distinguished through clonal data, once some time has passed after induction of clones. For GIA models, distributions can generally vary, but if the rates of divisions and transitions in the ℛ compartment are much larger that the rate of cell loss, the clone size distribution of all those models becomes a Normal distribution. In that case, two GIA models can not be distinguished by the clonal data. Hence, our analysis shows that models of cell fate choice cannot in general be distinguished with further resolution beyond the *R* vs. *C* categorization of cell types. The universality of the model dynamics also shows that effective, simplistic models are often equally accurate to model experimental data, yet with a higher statistical power due to less free parameters.

While at first glance this analysis seems to discourage efforts to unravel details of cell fate dynamics, room remains in regimes where the limiting conditions for asymptotic distributions are not fulfilled. In particular, if fast cycling committed progenitor cells are present, while stem cells are slow cycling, then the condition that the division rate of *R*-cells is much larger than the cell loss rate is not fulfilled. In that case, details of the model dynamics may affect the shape of the clone size distribution and thus allow distinction between models. However, caution should be given when an Exponential clone size distribution is observed, since this could indicate either a GIA model with high activity of committed progenitor cells, or a GPA model. In that case, the mean clone size needs to be consulted to distinguish models (see Fig. 2). Differentiating between models within the GPA category is more difficult, since the predicted statistics from different models always become more similar over time. Short term measurements would in principle allow such a distinction, but since in reality the underlying processes are not truly Markovian (as assumed for the modelling purpose) they are not necessarily a good representation of the real cell dynamics at short times. At long times however, Markovian approximations are increasingly accurate, precisly because of the feature of universality.

How could the resolution of cell fate modelling be improved? The state-of-the-art approach to determine cell fate trajectories is via analysis and modelling of single-cell RNA-sequencing (scRNA-seq) data. However, many limitations to this method exist, discussed in Ref. (29), and neither reversible trajectories nor the modes of cell division, such as asymmetric vs symmetric divisions, can be inferred. Intravital live imaging, on the other hand, allows to trace individual clones over time (30–33), and thus can obtain details of cell fate trajectories, yet this technique is limited to few tissue types which are accessible for invasive long-term imaging. Nonetheless, while each of those experimental assays alone is prone to limitations in defining self-renewal strategies, advanced model inference schemes, that integrate data from different experimental sources, might be the way forward in the future to finally reveal the details of stem cell self-renewal strategies.

## Materials and Methods

The numerical analysis of the random cell fate model was implemented in Matlab. The description of the stochastic models definition, the random model generation and the simulation campaign is detailed in the SI, section 2. Additionally, as a validation of the implemented simulator, based on the Gillespie algorithm (27), the IA and PA models were simulated and the results analyzed in the SI, section 3.A.

Analytical solutions were partially obtained using Mathematica.

## ACKNOWLEDGMENTS

We thank Benjamin D. MacArthur for valuable discussions that contributed in the development of this research. C.P. is supported by a Studentship of the Institute for Life Sciences (Southampton) and P.G. by Medical Research Council New Investigator Research Grant MR/R026610/1.

The authors declare no conflict of interest.

## Footnotes

∗ We note that we consider here stability in the sense of Lyapunov. Since the system is linear, the only asymptotically stable state is 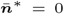 which does not correspond to a biological relevant homeostatic state. For a further discussion, see the SI section 1.

† The Perron-Frobenius theorem assures that for adjacency matrices of SCCs of cooperative systems, a unique, real, maximal eigenvalue exists, which is the dominant eigenvalue (25, 26).

‡ Since *µ* = 0 is necessary for conservation, the only possible conserved cell states in homeostasis are those in ℛ

§ In one dimension, i.e. below the critical dimension interactions between cells can significantly alter clonal statistics (5, 23), which is not considered here.

