## Supplemental Information (SI) for "Universality of clonal dynamics poses fundamental limits to identify stem cell self-renewal strategies"

### 1. Conditions for homeostasis

Here we "translate" the generic conditions for the existence of a Lyapunov stable stationary state for Linear Cooperative Systems ($LCS$) (1) into the biological context of clonal dynamics. A linear cooperative system is one of the form $\frac{d}{dt}\boldsymbol{x}(t) = A\boldsymbol{x}(t)$ where $\boldsymbol{x}(t) = (x_1(t), x_2(t), ..., x_m(t))$ are functions of time $t$ and $A$ is a constant $m \times m$ matrix for which all off-diagonal elements are non-negative (the latter condition defines the cooperativity of the system) (1, 2). We note that the dynamics of mean cell numbers, Eq. 6 and 7 in the main text, indeed describe an LCS according to this definition. Now we use the following definitions:

- $G(A)$ is the graph of $A$, i.e. the graph for which $A$ is the adjacency matrix, whose elements $a_{ij}$ give the weight of the links from $i$ to $j$ ($a_{ij} = 0$ means that no link exists). In the following we use the terms *graph* and *network* synonymously.
- If in $G(A)$ there exists a path from node $i$ to node $j$ *and* from $j$ to $i$, then we call those nodes *strongly connected*, $i \equiv j$, which is an equivalence relation. A maximal set of nodes which are are strongly connected with each other are called a *Strongly Connected Component (SCC)* of the graph (the equivalence class of the equivalence relation "$\equiv$").
- The graph $G(A)$ can be decomposed into its $N_S$ SCCs, $S_k$, $k = 1, ..., N_S$ (3), which are sub-graphs associated with an adjacency matrix $A_k$, such that $G(A_k) = S_k$. Since the $A_k$ have non-negative off-diagonal elements, they are Metzler matrices for which the Perron-Frobenius theorem ensures that a unique, simple and real maximal eigenvalue $\mu_k$ exists (4). The eigenvalue $\mu_k$ is called the *dominant eigenvalue* of $S_k$. Associated with this eigenvalue, there is, for all $k$, a positive eigenvector $\boldsymbol{x}^{(k)} = (x_1^{(k)}, x_2^{(k)}, ...)$, i.e. one with all entries $x_i^{(k)} > 0$.
- The *condensed graph* of $G(A)$ is the graph where nodes are the SCCs of $G(A)$ and a link from SCC $S_k$ to SCC $S_l$ ($k, l = 1, ..., N_S$) exists if there is is at least one link from a node (in $G(A)$) in $S_k$ to a node in $S_l$.
- If there is a path from SCC $S_k$ to SCC $S_l$, then we call $S_k$ *upstream* of $S_l$ and accordingly $S_l$ *downstream* of $S_k$. We note that there can never exist paths from $S_k$ to $S_l$ **and** from $S_l$ to $S_k$, since otherwise, by definition, their nodes would be strongly connected and both together would form a single SCC (3). Thus, there is a unique hierarchy of SCCs.
- A stationary state $\boldsymbol{x}^*$ of a dynamical system is *Lyapunov stable* if a small initial deviation from $\boldsymbol{x}^*$ leads to a small final deviation $x(t)$ (i.e. $\boldsymbol{x}^*$ is not unstable). More accurately: there exists a constant $C > 0$ such that $|\boldsymbol{x}(t) - \boldsymbol{x}^*| < C|\boldsymbol{x}_0 - \boldsymbol{x}^*|$ for all times $t$, where $\boldsymbol{x}_0 = \boldsymbol{x}(t = t_0)$ is the initial condition, sufficiently close to $\boldsymbol{x}^*$. A stationary state of a linear system that is Lyapunov stable, yet neither asymptotically stable nor has a limit cycle, is *neutrally stable*.
- Homeostasis means that the cell numbers in each state, $\boldsymbol{n} = (n_1, ..., n_m)$, stay on average constant, $\frac{d\bar{\boldsymbol{n}}}{dt} = 0$ (where $\bar{\boldsymbol{n}} = \langle \boldsymbol{n} \rangle$), and that this state is not unstable towards perturbations. This condition corresponds to a Lyapunov-stable stationary state. Note that a linear system, as the one described by Eqs. 6 and 7, main text, cannot have an asymptotically stable state except for the trivial state $\bar{\boldsymbol{n}}^* = 0$, which corresponds to a vanishing cell population*. We therefore use Lyapunov stability, a weaker form of stability, to define homeostasis, since an asymptotically stable vanishing state is not a biologically viable state.

*We note that when considering the tissue cell population as a whole, dynamics can be non-linear through interactions between cells and a non-vanishing asymptotically stable state may then exist. However, since single clones do not significantly affect the total configuration of cells in a tissue, the clones compete neutrally, when embedded in a homeostatic cell population, which corresponds to a Lyapunov stable, but not asymptotically stable state.

Now, for an LCS holds, according to Ref. (1),

**Theorem 1** *An LCS, $\dot{\boldsymbol{x}} = A\boldsymbol{x}$, possesses a non-trivial Lyapunov stable stationary state ($\boldsymbol{x}^* > 0$), if and only if,*

1. *$G(A)$ does not contain any SCC, $S_k$, with $\mu_k > 0$.*
2. *There is at least one SCC, $S_k$, with $\mu_k = 0$.*
3. *There is no path between any two SCCs, $S_k$ and $S_l$, which have $\mu_k = 0$ and $\mu_l = 0$.*

Furthermore holds,

**Theorem 2** *all nodes $i$ upstream of an SCC $S_l$ with $\mu_l = 0$ must be empty in the the stationary state, i.e. $x_i^* = 0$, if $i$ is upstream of the SCC $S_l$.*

Since Eq. 7, main text, is an LCS, we can apply theorems 1 and 2 to find conditions for homeostasis, defined by a Lyapunov-stable configuration of mean cell numbers $\bar{\boldsymbol{n}}^* = (\bar{n}_1, \bar{n}_2, ...)$. According to theorem 1 at least one SCC with $\mu_k = 0$ must then exist, and according to theorem 2 the stationary state of nodes upstream of it must be empty, i.e. they do not exist in homeostasis. Since the condensed graph of the SCCs does not have cyclic paths, an SCC $S_k$ with $\mu_k = 0$ must therefore always reside at the apex of all non-vanishing cell types. In principle, an acyclic graph may have more than one apex, however, since, by definition, a stem cell clone always starts with a single stem cell, and no other SCC with $\mu = 0$ may be downstream of the latter, we only consider one apex SCC with one initial cell when studying clonal dynamics.

Hence, in the context of homeostatic clonal dynamics, we can assume that there is a single SCC, $S_k$ with $\mu_k = 0$ at the apex of the cell state graph, while all other SCCs, $S_l$ are downstream of it and have $\mu_l < 0$. Since there are no paths from the non-apex SCC to the apex SCC (as the condensed graph is acyclic) we can distinguish the two separate compartments $\mathcal{R}$ (the renewing compartment) consisting of all nodes of the apex SCC, $S_k$, and $\mathcal{C}$ (the committed compartment), consisting of all other nodes, whereby due to $\mu_l < 0$ for all SCCs in $\mathcal{C}$, all progeny of cells in $\mathcal{C}$ will vanish in the long term.

### 2. Stochastic process modelling

**A. Model Description.** Since clonal dynamics start, by definition, with a single cell, we use stochastic dynamics to model clones. Thus, we model cell fate dynamics as a continuous-time multi-type branching process (5), a Markov process following the rules of Eq. 3-5, main text. As shown later, without losing generality, here only two types of events are modelled; considering an arbitrary number $m$ of cell states, $X_i$, for $i = 1, ...m$, the model includes

- **Cell divisions**: a cell in state $X_i$ divides in two cells respectively in state $X_j$ and $X_k$ at a given rate $\lambda_i$.

$$X_i \xrightarrow{\lambda_i} X_j + X_k, \qquad i, j, k = 1, ..., m, \tag{1}$$

  where $\lambda_i = 0$ if state $X_i$ does not allow division. In this formulation of cell division events, which we use for the generation and numerical simulations of random models, only one division outcome is possible upon division of a particular cell state $X_i$. Nonetheless, multiple division outcomes per state can be implemented as single outcomes if additional *metastates* are introduced, which represent priming of a state $X_i$ towards a certain division outcome option. For example, if in the original model, state $X_i$ has different outcome options, $X_{j_1} + X_{k_1}, X_{j_2} + X_{k_2}, ...$, we can substitute this by, first, transitions from $X_i$ to (new) states $X_{m_1}, X_{m_2}, ...$ and subsequent divisions $X_{m_l} \to X_{j_l} + X_{k_l}$. The use of metastates to model more complex processes is discussed in detail in Section 3.B.

- **Direct state transitions**: a cell in state $X_i$ changes to state $X_j$ at a given rate $\omega_{ij}$.

$$X_i \xrightarrow{\omega_{ij}} X_j, \qquad i, j = 1, ..., m;\ i \neq j, \tag{2}$$

  where $\omega_{ij} = 0$ means that no transition from $X_i$ to $X_j$ is possible. Additionally, we include cell loss in this scheme, by treating it as a transition to an additional special state, called hereafter *death* and denoted by $\emptyset$ (cells in this state do not enter in the counting of the total number of cells). In that formulation, the loss rates of the original model are $d_i = \omega_{i\emptyset}$.

These events define a Markov process, which can be represented as a *stochastic network* (6). In this view, each node can be related to a cell state, while the links represent transitions between states via cell divisions and the direct state transitions. It is noted that this stochastic network is different from the network defined in the main text and in section 1 of this SI, which describes the dynamics of *mean* cell number instead. Here, for the stochastic modelling, let us define the adjacency matrix $K$ of this network, through the elements $\kappa_{ij} = \lambda_i 2 r_i^j + \omega_{ij}$ $i, j = 1, ..., m$, in which $\kappa_{ij}$ are the total transition rates as defined in the main text. We note that $K$ is related to the matrix $A$ used in the main text by $A = K^T - \Delta$, where $\Delta$ is the diagonal matrix with entries $\delta_i, i = 1, ..., m$, as defined in the main text, with the slight difference that here the loss state $\emptyset$ is treated as a separate state. Additionally, it is remarked that in this model interpretation, where only one division option for each state is possible, the term $r_i^j \leq 1$ is not a continuum value, but instead it can only take the values $0, 1/2, 1$ depending on the specific outcome of the division of the cells in state $X_i$. Notably, more than one stochastic network may result in the same matrix $K$, therefore, to uniquely define a process, we distinguish a matrix $D$ which describes cell division events (note that this is possible with just a single matrix as there is only one division option per state) and a matrix $T$ which describes direct transition events. The matrix $K$ is the sum of both, $K = N + T$.

**B. Generation of Random Models.** To test the behaviour of the clonal dynamics in a generic homeostatic model, a large number of random stochastic networks was generated, whereby each stochastic network corresponds to a distinct set of parameters $\lambda_1, ..., \lambda_m, \omega_{12}, ..., \omega_{m\emptyset}$ for the stochastic stem cell fate choice model. The strategy detailed below is based on the following considerations which summarize the key requirements to achieve homeostasis detailed in section 1: a) each network is composed of Strongly Connected Components (SCCs) that are randomly connected; b) only one SCC, the one at the apex of the network, forms the renewing compartment, $\mathcal{R}$, (i.e. it is characterized by a dominant eigenvalue $\mu = 0$ with respect to $A$) and all the others form the committed compartment, $\mathcal{C}$, (i.e. they are characterized by a dominant eigenvalues $\mu < 0$). It is further noted that the SCCs of the stochastic network $G(K)$ are the same as those of the matrix $G(A)$, where $A = K^T - \Delta$ defines the dynamics of mean cell numbers. This is, since transposition of an adjacency matrix and altering of diagonal elements does not affect the network topology.

To generate the stochastic network, a two-step process is followed: 1) a large number of (random) SCCs are generated; 2) a condensed network is randomly constructed and filled with randomly picked SCC from step 1.

It is noted that unitary rates are assumed in step 1) and they are successively randomly modified in step 2) to achieve the desired properties of the dominant eigenvalue $\mu$ while ensuring randomness.

Focusing now on step 1), that is, the generation of single SCCs, the following procedure is used.

(1.a) The total number of states composing the SCC is defined, indicated as $m_S$. An additional state is added to represent whatever is outside the SCC. In the current analysis we set $1 \leq m_S \leq 4$.

(1.b) We build separately all the possible combinations of transition and division matrices, indicated hereafter respectively with $M_T$ and $M_D$. These matrices are ordered for increasing number of transitions $N_T$ and divisions $N_D$. In case GIA networks are generated, the $M_D$ and $M_T$ combinations are filtered, to remain just with those where the division outcome is one cell inside the SCC and one outside the SCC, and where there are only transitions between states within the SCC (i.e. where cell numbers are conserved). From a computational point of view, this process is feasible up to $m_S = 4$.

(1.c) The matrices stored in $M_D$ and $M_T$ are then combined together to form a model (which is completely defined by one matrix in $M_D$ and one in $M_T$); $M_{DT}$ indicates the pool of possible models. This process is done considering separately each $m_S$, $N_T$ and $N_D$. In this step, due to technical limitations given by the high number of possible combinations, if the total number of combinations exceed $5 \cdot 10^4$ then only $10^4$ random matrices from $M_D$ and $M_T$ are combined.

(1.d) Each model in $M_{DT}$ is then processed to check if the corresponding network is a SCC in the first $m_S$ states. If not, then this model is discarded. In case GPA networks are generated, a further check is performed to discard also those models consistent with a GIA network (they cannot be a priori excluded as done in point (1.b) for the GIA ones). These pools of models are indicated as $M_{\mathrm{GIA}}$ and $M_{\mathrm{GPA}}$ respectively for the GIA and GPA models.

(1.e) For each SCC in $M_{\mathrm{GIA}}$ and $M_{\mathrm{GPA}}$, the dominant eigenvalue $\mu$ is estimated. For construction, the generated GIA networks are all characterized by $\mu = 0$, while in general any value can be obtained within $M_{\mathrm{GPA}}$.

(1.f) The SCCs in $M_{\mathrm{GPA}}$ are additionally processed to check whether the network is compatible with homeostasis by tuning the rates. Networks satisfying this condition are additionally stored under a new pool of SCCs, called $M^*_{\mathrm{GPA}}$. If not, then they are discarded when $\mu > 0$ (i.e. for any combination of rates the number of cells in these networks is expected to grow).

This process results in three pools of SCCs classified for $m_S$, $N_T$ and $N_D$ (i.e. number of states, transitions and divisions): 1) $M_{\mathrm{GIA}}$ contains GIA models; 2) $M^*_{\mathrm{GPA}}$ contains GPA models that can be tuned to have $\mu = 0$ and 3) $M_{\mathrm{GPA}}$ contains GPA models characterized by $\mu < 0$ or that can be tuned to meet this condition.

In step 2), the generation of random networks starting from the individual SCCs is implemented as follows.

(2.a) A number of committed SCCs, $N_c$, between 1 and 3 is randomly chosen.

(2.b) $N_c$ SCCs are randomly picked from the pool of models $M_{\mathrm{GPA}}$. The selection is done considering equal probability in $m_S$, $N_T$ and $N_D$. For each SCC, the unitary rates $\alpha$ (where $\alpha$ stands for any rate $\lambda_i$ or $\omega_{ij}$) are modified by multiplying them for random numbers (exponentially distributed with mean $\bar{\alpha} = 1$ and minimum $\alpha_m = 0.3$). Additionally, a threshold on the dominant eigenvalue is set, $\mu_{\max} = -1$; if this condition is not satisfied, then the rates are tuned to meet this requirement while maintaining the rates above the minimum.

(2.c) The committed compartment of the condensed network is generated by randomly connecting all the outgoing components of the $k$-SCC with states in the $l$-SCC for $l = k+1, .., N_c$. In this way, the transposed adjacency matrix of the stochastic network has triangular block form:

$$K^T = \begin{bmatrix} B_1 & & & \\ C_{12} & B_2 & 0 & \\ & & ... & \\ C_{1,N_c} & C_{2,N_c} & B_{N_c} & \\ C_{1\emptyset} & C_{2\emptyset} & C_{N_c,\emptyset} & 0 \end{bmatrix}. \qquad [3]$$

The last SCC is forced to be linked to a single death state.

(2.d) With a similar procedure described in point (2.b), two SCCs are randomly picked respectively from the pool of SCCs in $M^*_{\mathrm{GPA}}$ and $M_{\mathrm{GIA}}$; the unitary rates are modified (exponentially distributed with mean $\bar{\alpha} = 1$ and minimum $\alpha_m = 0.3$) and, in the GPA case, tuned to meet the condition $\mu = 0$. They represent the renewing part of the network.

(2.e) Two networks (one for the GIA and one for the GPA models) are produced by attaching the selected renewing network upstream the committed one; this is done based on an analogous procedure as described in step (2.c).

At the end of this process we have two networks which are different in just the renewing part, being one consistent with the GIA model and the other with the GPA one. In total 2000 networks were built and analysed.

**C. Simulation campaign.** An extensive simulation campaign was run to model the clone dynamics. The code implemented to numerically simulate the stochastic process defined by events of type 1 and 2 is based on the Gillespie algorithm (7). Since a clone is by definition the progeny of a single cell, we choose as initial condition a single cell put randomly in a state within $\mathcal{R}$. Concerning the final condition, given the substantial difference in the dynamics in the two models, the final time, indicated by $\tau$, is set equal to 20 times the inverse of the minimum process rate, $\alpha_{\min} = \min(\lambda_1, ..., \lambda_m, \omega_{12}, ..., \omega_{m,\emptyset})$, in the GIA models, and to the time at which the fraction of extinct clones reaches 98% in the GPA models †.

To determine the clone size distribution, $10^3$ and $5 \cdot 10^4$ simulations were run respectively in for each GIA and GPA model (in this way, both models result in the same final number of clones when 98% extinction is taken into account).

### 3. Numerical Simulation Test Cases

**A. Invariant Asymmetry and Population Asymmetry Models.** To validate the simulation approach, we tested the procedure on simple cell fate models for which analytical results are known, the Invariant Asymmetry (IA) and Population Asymmetry (PA) models. As described in the main text, in the simplest version, these are defined as

$$S \xrightarrow{\lambda} \begin{cases} S+S & \text{Pr. } r \\ S+D & \text{Pr. } 1-2r \\ D+D & \text{Pr. } r \end{cases}, D \xrightarrow{\gamma} \emptyset. \quad [4]$$

In these processes, cells of type $S$ represent the stem cells (called hereafter also progenitor), which divide with stochastic rate $\lambda$, and cells of type $D$ are the differentiated cells, which are shed with rate $\gamma$. While in the PA model the three possible outcomes of the division of a progenitor are controlled by a probability parameter $0 < r \leq 1/2$, in the IA model $r = 0$, meaning that there are strictly asymmetric division and the number of $S$-cells is conserved. It is remarked that in the definition of the stochastic networks given in section 2.A only one division option for each state is modelled; however, the code implemented for the numerical simulations of the stochastic process allows for an arbitrary number of division options for each state as well (see section 3.B).

Considering the dynamics at tissue level, the system of ODEs describing the average number of cell $\bar{n}_S$ and $\bar{n}_D$ respectively of type $S$ and $D$ is

$$\begin{cases} \dfrac{d\bar{n}_S}{dt} = 0 \\ \dfrac{d\bar{n}_D}{dt} = \lambda\bar{n}_S - \gamma\bar{n}_D \end{cases}. \quad [5]$$

It is clear that, on average, the number of $S$-cells remains constant. Additionally, in homeostasis, the average total number of $D$-cells stabilizes around a constant value $\bar{n}^*_D = (\lambda/\gamma)\,\bar{n}_S$ that uniquely depends on the number of stem cells, $\bar{n}_S$ which equals the initial number of stem cells $\bar{n}_{S,0} = \bar{n}_S(t=0)$, Thus, the (Lyapunov stable) stationary state of total cell numbers $\bar{n} = \bar{n}_S + \bar{n}_D$ is given by

$$\bar{n}^* = \left(1 + \frac{\lambda}{\gamma}\right)\bar{n}_{S,0}. \quad [6]$$

Based on Eq. 6, the process rates $\lambda$ and $\gamma$ determine the proportion of cells of type $D$ with respect to cells of type $S$. Importantly, there is no difference at tissue level between the IA and PA models.

A distinction is instead evident when we look at the dynamics at the single cell level, and study the clone size distribution, that is, the distribution of the progeny of a single cell. For the IA model, the number of $S$-cells is strictly constant, and thus the joint probability distribution $P(n_S, n_D)$ of both $S$-cells and $D$-cells, respectively indicated as $n_S$ and $n_D$, is fully determined by the distribution of $D$-cells, $P(n_D)$. The IA model's master equation for $P(n_D)$, considering a single initial cell of type $S$, is given by,

$$\frac{dP(n_D)}{dt} = \lambda P(n_D - 1) + \gamma(n_D + 1)P(n_D + 1) - (\lambda + \gamma n_D)\,P(n_D). \quad [7]$$

This corresponds to a simple birth-and-death process for which the distribution is Poissonian with mean $\lambda/\gamma$, (8).

† Note that all critical branching processes, as homeostatic clonal dynamics are, will go extinct almost surely at some point in time (5).

Considering now the PA model, the master equation is instead given by

$$\begin{aligned}\frac{dP(n_S,n_D)}{dt} =&\lambda(r(n_S-1)P(n_S-1,n_D)+(1-2r)n_SP(n_S,n_D-1)+r(n_S+1)P(n_S+1,n_D-2))\\ &+\gamma(n_D+1)P(n_S,n_D+1)\\ &-(\lambda n_S+\gamma n_D)\,P(n_S,n_D).\end{aligned} \quad [8]$$

In Ref. (9), an exact result for the distribution of total cell numbers $n = n_S + n_D$ is found when $\lambda = \gamma$ and $r = 1/4$. For different values of the process parameters, the long-term distribution is shown to be Exponential.

Numerical simulations for the clonal dynamics were run, considering the above models and three different sets of test parameters each, indicated as IA$\#i$ and PA$\#i$ for $i = 1, 2, 3$, which are reported in Table 1. It is noted that the time unit is arbitrary and therefore omitted. Simulations are based on $10^4$ and $5 \cdot 10^4$ runs respectively for the IA and PA test cases. The initial condition is a single stem cell and the final simulation time, indicated as $\tau$, is equal to 10: this value is well representative of a steady state condition (for the IA test cases) and at which the total extinction of the process is not yet achieved (for PA test cases only). The clone size distribution at $\tau$ in the IA test cases is shown in Figure 1: in this figure, each profile is compared to the corresponding Poisson distribution shifted by one (i.e. plus the stem cell). Concerning the results for the PA test cases, they are shown in Figure 2. In this case, the profiles are compared to the numerical integration of the master equation 8. Additionally, for the PA#1 test case, where $\lambda = \gamma$ and $r = 1/4$, the reference analytic solution provided in Ref. (9) is also shown. In general, a good agreement is obtained in all of the cases.

**B. Population Asymmetry Model Using Metastates.** As argued before, we assume in the random model generation that cell division in state $X_i$ has a unique outcome, $X_i \to X_j + X_k$ (Eq. 1), since thereby the stochastic process can be uniquely defined by the two matrices $D$ and $T$. To accommodate for the possibility of different division outcomes from the same state $X_i$, as in Eq. 4 and Eqs. 3-5 in the main text, we introduce *metastates*, which represent short-lived states that indicate priming for either outcome, from which the cell division outcomes are unique. This is a small modification of the original model, which, however, does not lead to significant deviations if the metastates are traversed sufficiently quickly (which can be assured by a choice of high direct state transition rates in the metastates).

To illustrate this, let us consider the PA model described by 4; instead of having three different outcomes upon division of an S-cell we define the corresponding Metastate (MS) model with three primed states, $M_{1,2,3}$, as

$$\begin{aligned} &S \xrightarrow{\omega_1} M_1,\ M_1 \xrightarrow{\lambda_1} S+S,\\ &S \xrightarrow{\omega_2} M_2,\ M_2 \xrightarrow{\lambda_2} S+D,\\ &S \xrightarrow{\omega_3} M_3,\ M_3 \xrightarrow{\lambda_3} D+D,\\ &D \xrightarrow{\gamma} \emptyset,\end{aligned} \quad [9]$$

in which $S$ and $D$ correspond to the same cell type of the PA model (i.e. respectively the stem and the differentiated cells), while $M_i$, for $i = 1, 2, 3$, represent the metastates. These states are temporary states that are used to model each one of the three different possible division options of the $S$-cells. The rates $\lambda_i$ and $\omega_i$, for $i = 1, 2, 3$, are chosen such that the time scales of division and outcome probabilities are the same as in the original PA model:

$$\omega_1/\omega_2 = r/(1-2r),\ \omega_2/\omega_3 = (1-2r)/r, \quad [10]$$

$$\frac{1}{(1/\omega_1+1/\lambda_1)} = \lambda r,\ \frac{1}{(1/\omega_2+1/\lambda_2)} = \lambda(1-2r),\ \frac{1}{(1/\omega_3+1/\lambda_3)} = \lambda r. \quad [11]$$

Eqs. 10 assure that outcome probabilities are the same as in the original model, while Eqs. 11 are needed to have the same total average time between two consecutive events. As there are six unknowns and only five relations, the following additional equation is added

$$\lambda_1 = \omega_1\Delta, \quad [12]$$

in which $\Delta$ is an additional parameter that is used to control how fast cells in metastate $M_1$ divide. Low values of $\Delta$ imply that as soon as an $S$-cell transits to the metastate $M_1$, it divides in two $S$-cells. Globally, this results in

$$\begin{aligned}\omega_1 &= \omega_3 = \lambda r(\Delta+1)/\Delta\\ \omega_2 &= \lambda(1-2r)(\Delta+1)/\Delta.\\ \lambda_i &= \omega_i\Delta \text{ for } i=1,2,3\end{aligned} \quad [13]$$

Numerical simulations for the two models were run and compared, based on the parameters reported in Table 1, and specifically the PA#1 and PA#3 test cases. The time unit, which is arbitrary, is omitted. The process rates for the corresponding MS model, which are indicated in the figures as MS#1 and MS#3, are computed based on Eq. 13 and $\Delta = 1/500$. As well as for the PA test cases, the initial condition is one cell of type $S$ and the final time, $\tau$, is equal to 10; simulations are based on $5 \cdot 10^4$ trajectories.

The mean number of cells in the surviving clones and the extinction probability as function of time (scaled by $\tau$) are shown in Figure 3. The clone size distribution at $\tau$ is shown in Figure 4. Both MS simulations agree very well with the corresponding PA ones, which justifies the use of metastates for our simulation campaign.

### 4. Analysis of the Generalized Invariant Asymmetry Model

**A. GIA$^0$ Test Case: Steady State Distribution and Limiting Behaviour.** A simple Generalized Invariant Asymmetric model, indicated hereafter as GIA$^0$, was analyzed to identify the causes of the different clone size distribution behaviours observed in the randomly generated models (see main text). Thus, in this section, we study the Markov process defined by

$$X_1 \xrightarrow{\lambda_1} X_1 + X_2, X_2 \xrightarrow{\lambda_2} X_2 + X_2, X_2 \xrightarrow{\gamma} \emptyset. \tag{14}$$

Here, the renewing compartment is composed of just a single state $X_1$ and cells in this state asymmetrically divide with rate $\lambda_1$. The committed compartment is formed of state $X_2$; cells in this state can either divide to duplicate, with rate $\lambda_2$, or die, with rate $\gamma$. It is noted that for $\lambda_2 = 0$, this model is reduced to the previously analyzed Invariant Asymmetric (IA) model (see section 3.A).

As for the IA model, here the number of cells in state $X_1$, indicated as $n_1$, is conserved. It is therefore sufficient to determine the statistics of $n_2$, defined by the master equation for $P(n_2)$, the probability of having $n_2$ cells in state $X_2$, provided that there are $n_1$ cells in state $X_1$. The master equation is given by

$$\begin{aligned} \frac{dP(n_2)}{dt} = &- (\lambda_1 n_1 + \lambda_2 n_2 + \gamma n_2)\, P(n_2) \\ &+ (\lambda_1 n_1 + \lambda_2 (n_2 - 1))\, P(n_2 - 1) \\ &+ \gamma (n_2 + 1) P(n_2 + 1), \end{aligned} \tag{15}$$

also written as

$$\begin{aligned} \frac{dP(n_2)}{dt} = &- (g(n_2) + r(n_2))\, P(n_2) \\ &+ g(n_2 - 1)P(n_2 - 1) + r(n_2 + 1)P(n_2 + 1), \end{aligned} \tag{16}$$

in which $r(n_2) = \gamma n_2$ and $g(n_2) = \lambda_1 n_1 + \lambda_2 n_2$. Considering that we are interested in clonal dynamics, meaning that we start from a single stem cell, $n_1$ is equal to one.

In this simple case, the steady state distribution $P^*(n_2)$, corresponding to the solution of $dP(n_2)/dt = 0$, can be analytically derived. Defining the net flux between states $n_2$ and $n_2 - 1$ as

$$I_{n_2} = r(n_2)P^*(n_2) - g(n_2 - 1)P^*(n_2 - 1), \tag{17}$$

and considering that $I_{n_2+1} = I_{n_2}$ for every $n_2$, it follows that $I_{n_2} = I_0 = r(0)P^*(0) - g(-1)P^*(-1) = 0$, which means that

$$P^*(n_2) = \frac{g(n_2 - 1)}{r(n_2)} P^*(n_2 - 1) = \prod_{l=0}^{n_2 - 1} \frac{g(l)}{r(l+1)} P^*(0), \tag{18}$$

where $P^*(0)$ is the steady state probability of having 0 cells in state $X_2$. Finally, by applying the conservation of the total probability, $\sum_{n_2=0}^{\infty} P^*(n_2) = 1$, and rearranging the terms we obtain

$$P^*(n_2) = \left(1 - \frac{\lambda_2}{\gamma}\right)^{\lambda_1/\lambda_2} \left(\frac{\lambda_2}{\gamma}\right)^{n_2} \frac{\Gamma\left(\frac{\lambda_1}{\lambda_2} + n_2\right)}{\Gamma(n_2 + 1)\Gamma\left(\frac{\lambda_1}{\lambda_2}\right)}. \tag{19}$$

In the main text we defined the dimensionless parameters $\hat{\lambda}_1 = \lambda_1/\gamma$ and $\hat{\lambda}_2 = \lambda_2/\gamma$, representing the rescaled division rates respectively for cells in state $X_1$ and $X_2$. For clarity and readability, in this section, we simplify the notation using $p = \hat{\lambda}_1$ and $q = \hat{\lambda}_2$. Eq. 19 is then rewritten as

$$P^*(n_2) = (1 - q)^{p/q}\, q^{n_2} \frac{\Gamma\left(\frac{p}{q} + n_2\right)}{\Gamma(n_2 + 1)\Gamma\left(\frac{p}{q}\right)}. \tag{20}$$

It is noted that while $p$ varies between 0 and $\infty$, $q$ is defined between 0 and 1.

The mean number of cells in each state, indicated respectively as $\bar{n}_1$ and $\bar{n}_2$, satisfies the system of ODEs

$$\begin{cases} \dfrac{d\bar{n}_1}{dt} = 0 \\ \dfrac{d\bar{n}_2}{dt} = \lambda_1 \bar{n}_1 + (\lambda_2 - \gamma)\bar{n}_2 \end{cases}. \tag{21}$$

Based on this, the steady state average number of cells is

$$\begin{cases} \bar{n}_1^* = 1 \\ \bar{n}_2^* = \dfrac{\lambda_1}{\gamma - \lambda_2} = \dfrac{p}{1 - q} \end{cases}. \tag{22}$$

When the mean number of cells in state $X_2$ is sufficiently large, i.e. for large $p$ or in case $q$ is close to one, the discrete distribution given by equation 20, can be approximated by a continuous probability density function $P^*(x_2)$, given by

$$P^*(x_2) = (1-q)^{p/q} q^{px_2/(1-q)} \frac{\Gamma\left(\frac{p}{q} + \frac{p}{1-q}x_2\right)}{x_2 \Gamma\left(\frac{p}{q}\right)\Gamma\left(\frac{p}{1-q}x_2\right)}, \quad [23]$$

in which $x_2 = n_2/\bar{n}_2^*$. We note that Eq. 23 corresponds to Eq. 11 in the main text.

To better understand the distribution for different values of the parameters $p$ and $q$, the limit behaviour are analysed below.

1. $\mathbf{q \to 0}$ (i.e. $\hat{\lambda}_2 \to 0$)

   When $q \to 0$, Eq. 20 can be simplified considering that

$$\lim_{q\to 0} \frac{\Gamma\left(\frac{p}{q} + n_2\right)}{\Gamma\left(\frac{p}{q}\right)} \left(\frac{q}{p}\right)^{n_2} = 1, \quad [24]$$

$$\lim_{q\to 0} (1-q)^{p/q} = e^{-p} \quad [25]$$

   and

$$\Gamma(n_2 + 1) = n_2!. \quad [26]$$

   Thus, the distribution results in

$$\lim_{q\to 0} P^*(n_2) = \frac{p^{n_2} e^{-p}}{n_2!} = \text{Poisson}(p), \quad [27]$$

   that is a Poisson distribution with mean equal to $p$. This agrees with what we were expecting considering that when $q = 0$ the model is reduced to the IA model for which the distribution in $n_2$ is known to be poissonian.

   Additionally, for large mean number of cells, which are obtained for large $p$ (when $q = 0$, then $\bar{n}_2^* = p$), the Poisson distribution tends to a Normal distribution with mean and variance equal to $p$. Therefore,

$$\lim_{(q,p)\to(0,\infty)} P^*(n_2) = \frac{1}{\sqrt{2\pi p}} e^{-\frac{(n_2-p)^2}{2p}} = \text{Normal}(p, p). \quad [28]$$

   Rescaling the distribution, and considering $x_2 = n_2/\bar{n}_2^*$, results in

$$\lim_{(q,p)\to(0,\infty)} P^*(x_2) = \text{Normal}(1, 1/p), \quad [29]$$

   that is a Normal distribution with unitary mean and variance equal to $1/p$.

2. $\mathbf{q \to 1}$ (i.e. $\hat{\lambda}_2 \to 1$)

   For $q \to 1$ the steady state mean number of cells $\bar{n}_2^* \to \infty$ and Eq. 23 holds. This equation can be rewritten as

$$P^*(x_2) = q^{p/(1-q)x_2+1} \frac{(1-q)^{p/q}}{q(x_2-1)+1} \frac{\Gamma\left(p\frac{q(x_2-1)+1}{q(1-q)} + 1\right)}{\Gamma\left(\frac{p}{q}\right)\Gamma\left(\frac{p}{1-q}x_2 + 1\right)}. \quad [30]$$

   If the Stirling's approximation is applied

$$\Gamma(z+1) = \sqrt{2\pi z}\left(\frac{z}{e}\right)^z, \quad [31]$$

   we obtain

$$P^*(x_2) = \frac{p^{p/q} e^{-p/q} q^{(q-2p)/(2q)} (q(x_2-1)+1)^{p/(1-q)(x_2-1+1/q)-1/2}}{\Gamma\left(\frac{p}{q}\right) x_2^{x_2 p/(1-q)+1/2}}. \quad [32]$$

   Considering now that

$$\lim_{q\to 1} \frac{(q(x_2-1)+1)^{p/(1-q)(x_2-1+1/q)-1/2}}{x_2^{x_2 p/(1-q)+1/2}} = e^{p(1-x_2)} x_2^{p-1}, \quad [33]$$

it follows that

$$\lim_{q\to 1} P^*(x_2) = \frac{p^p}{\Gamma(p)} x_2^{p-1} e^{-px_2} = \text{Gamma}(p, 1/p), \quad [34]$$

that is a Gamma distribution with unitary mean and shape parameter given by $p$. Importantly, the Gamma distribution for $p \to \infty$ tends to a Normal distribution with unitary mean and variance $1/p$. For $p = 1$, it corresponds instead to an Exponential distribution with unitary mean.

3. $\mathbf{p} \to \infty$ (i.e. $\hat{\lambda}_1 \to \infty$)

When $p$ is large, the mean number of cells is large for any value of $q$. Thus, Eq. 32 is valid. By applying the Stirling's approximation also to the term $\Gamma(p/q)$, we obtain

$$P^*(x_2) = \sqrt{\frac{p}{2\pi}} x_2^{-p/(1-q)x_2 - 1/2} (q(x_2 - 1) + 1)^{p/(1-q)(x_2 - 1 + 1/q) - 1/2}. \quad [35]$$

This expression can be also rewritten as

$$P^*(x_2) = \sqrt{\frac{p}{2\pi}} e^{p/(1-q)((x_2-1+1/q)\log(q(x_2-1)+1) - x_2\log(x_2)) - 1/2(\log(x_2)+\log(q(x_2-1)+1))}. \quad [36]$$

Considering now that $p$ is large, then $-1/2(\log(x_2) + \log(q(x_2 - 1) + 1)) \ll p/(1 - q)((x_2 - 1 + 1/q)\log(q(x_2 - 1) + 1) - x_2 \log(x_2))$, so the term on the right can be neglected. Additionally, for $x_2 \to 1$ the following expansions can be applied

$$\log(q(x_2 - 1) + 1) = \sum_{k=1}^{\infty} \left( (-1)^{k+1} \frac{(q(x_2 - 1))^k}{k} \right), \quad [37]$$

and

$$\log(x_2) = \sum_{k=1}^{\infty} \left( (-1)^{k+1} \frac{(x_2 - 1)^k}{k} \right). \quad [38]$$

Finally if we consider that

$$\frac{\left(x_2 - 1 + \frac{1}{q}\right) \sum_{k=1}^{\infty} \left( (-1)^{k+1} \frac{(q(x_2-1))^k}{k} \right) - x_2 \sum_{k=1}^{\infty} \left( (-1)^{k+1} \frac{(x_2-1)^k}{k} \right)}{(x_2 - 1)^2} = -\frac{1}{2(1 - q)}, \quad [39]$$

then Eq. 36 results in

$$\lim_{p\to\infty} P^*(x_2) \simeq \sqrt{\frac{p}{2\pi}} e^{-1/2p(x_2-1)^2} = \text{Normal}(1, 1/p), \quad [40]$$

that is a Normal distribution with unitary mean and variance equal to $1/p$.

Importantly, it is noted that the limiting behaviour of $P^*(x_2)$ for $q \to 0$ and $q \to 1$ in case of large $p$, are both consistent with the results obtained for $p \to \infty$ and any $q$. In other words, remembering that $p = \hat{\lambda}_1$ and $q = \hat{\lambda}_2$, the steady state distribution for $\hat{\lambda}_1 \to \infty$ and any value of $\hat{\lambda}_2$ is a Normal distribution of unitary mean and variance equal to $1/\hat{\lambda}_1$.

To globally verify these results, numerical simulations of the stochastic process associated with model 14 for different values of $\hat{\lambda}_1$ and $\hat{\lambda}_2$ were run. The following curves were compared:

- **Stochastic simulation:** distribution at the final simulation time, $\tau$, of the number of cells in state $X_2$. The final time was chosen here as $\tau = 20/\alpha_{\min}$, where $\alpha_{\min} = \min(\lambda_1, \lambda_2, \gamma)$; this value is well representative of a steady state condition. Furthermore, the process rates considered are based on a unitary $\gamma$ (i.e. $\lambda_1 = \hat{\lambda}_1$, $\lambda_2 = \hat{\lambda}_2$ and $\gamma = 1$). It is noted that the time unit is arbitrary and therefore omitted.

- **Analytic distribution:** based on Eqs. 20, for low mean values, and 23, for large mean values.

- **Approximate distributions:** Poisson, Gamma and Normal distributions respectively given by Eqs. 27, 34 and 40.

The tested parameters $\hat{\lambda}_1$ and $\hat{\lambda}_2$ are graphically shown in Figure 5 a contour map showing the expected steady state mean number of cells $\bar{n}_2^*$ over the $(\hat{\lambda}_1, \hat{\lambda}_2)$-parameter plane. The curves from the numerical simulations and the corresponding exact and approximated solutions are shown in Figure 6, Figure 7 and Figure 8: the tested conditions are divided into three groups (one figure each) representing the limiting behaviours discussed above. Generally, analytical and numerical results agree very well. This also demonstrates that GIA models can show both peaked and non-peaked distributions, depending on the model parameters.

**B. Approximation of generic GIA Models.** As shown in the main text, a generic GIA model can be expressed in terms of the compartments $\mathcal{R}$ and $\mathcal{C}$ (Eq. 9 in the main text). We note that the the $\text{GIA}^0$ model discussed in the previous section corresponds to the general compartment dynamics of GIA models, Eq. 9, main text, if the dynamics of compartments are assumed to be Markovian. Thus, we can treat the $\text{GIA}^0$ model as a Markovian approximation of generic GIA models. In this section, we test this approximation numerically.

To this end, we first wish to relate the effective (non-Markovian) rates $\lambda_{R,C}$ and $\gamma_C$ of a generic GIA model to the rates of the Markovian approximation, the $\text{GIA}^0$ model. We refer to this model – the $\text{GIA}^0$ model matched to the effective rates of a particular more complex GIA model – as the *equivalent model* to the latter. The equivalent rates $\lambda_R$, $\lambda_C$ and $\gamma_C$ are computed considering the same steady state condition in terms of mean number of cells. To this aim, we rewrite the dynamics of mean cell numbers, Eq. 7 in the main text, in block form as

$$\begin{cases} \dfrac{d\bar{\mathbf{n}}_R}{dt} = A_{RR}\bar{\mathbf{n}}_R \\ \dfrac{d\bar{\mathbf{n}}_C}{dt} = A_{CR}\bar{\mathbf{n}}_R + A_{CC}\bar{\mathbf{n}}_C \\ \dfrac{d\bar{n}_\emptyset}{dt} = A_{\emptyset C}\bar{\mathbf{n}}_C \end{cases}, \tag{41}$$

in which $\bar{\boldsymbol{n}}_{R,C}$ denote the vectors of mean cell numbers of states restricted to compartments $\mathcal{R},\mathcal{C}$, respectively, and $n_\emptyset$ the number of lost cells (not considered for total cell numbers and homeostasis condition). It is noted that $A_{RC} = \mathbf{0}$, since there cannot be links from $\mathcal{C}$ to $\mathcal{R}$. Also $A_{\emptyset R} = \mathbf{0}$ as we do not consider loss from $\mathcal{R}$ (see main text for the arguments).

Thus, summing up all the components in each compartment, $\bar{n}_R = \sum_i (\bar{\mathbf{n}}_R)_i = 1$ and $\bar{n}_C = \sum_i (\bar{\mathbf{n}}_C)_i$, results in

$$\begin{cases} \dfrac{d\bar{n}_R}{dt} = 0 \\ \dfrac{d\bar{n}_C}{dt} = \sum_i (A_{CR}\bar{\mathbf{n}}_R)_i + \sum_i (A_{CC}\bar{\mathbf{n}}_C)_i \\ \dfrac{d\bar{n}_\emptyset}{dt} = A_{\emptyset C}\bar{\mathbf{n}}_C \end{cases}. \tag{42}$$

The equivalent parameters are then estimated from the steady state condition $\bar{\mathbf{n}}_X^*$ and $\bar{n}_X^*$, for $X = R, C, \emptyset$, as

$$\lambda_R = \sum_i (A_{CR}\bar{\mathbf{n}}_R^*)_i,\ \gamma_C = \frac{\sum_i (A_{\emptyset C}\bar{\mathbf{n}}_C^*)_i}{\bar{n}_C^*} \text{ and } \lambda_C = \gamma_C - \frac{\lambda_R}{\bar{n}_C^*}. \tag{43}$$

The applicability of this approximation was evaluated by comparing the clone size distribution obtained from the random GIA models (generated as described in 2.B and analyzed in the main text) with that from the corresponding equivalent $\text{GIA}^0$ model with parameters $\hat{\lambda}_1 = \hat{\lambda}_R = \lambda_R/\gamma_C$ and $\hat{\lambda}_2 = \hat{\lambda}_C = \lambda_C/\gamma_C$. The values of $\hat{\lambda}_1$ and $\hat{\lambda}_2$ for all the GIA random models are shown in Figure 9 in the contour map of the expected mean number of cells in $\mathcal{C}$ (in compartment $\mathcal{R}$ there is always one single cell). In general, $\hat{\lambda}_1$ remains below five and $\hat{\lambda}_2$ is spread between zero and one. As measure of the error of the equivalent model, $\epsilon$, we choose the maximum difference between the distributions of a particular random GIA model and that of the corresponding equivalent model, relative to the peak of the distribution of the random model. For low mean cell numbers, the distribution is compared to Eq. 20; for large mean number instead, the rescaled distribution is compared to Eq. 23. A threshold on the mean cell number equal to 10 was chosen to distinguish between the two cases. This relative error $\epsilon$ as function of $\hat{\lambda}_2$ is presented in Figure 10, where it is evident that large errors are obtained only for large values of this parameters. Some illustrative cases, representative of different value of $\hat{\lambda}_2$, were selected and their distribution is shown in Figure 11, Figure 12 and Figure 13. The following considerations are made:

- Two cases for $\hat{\lambda}_2 < 0.2$ are presented in 11. In these cases, the distribution obtained from the random models agrees with the analytic solution from the equivalent model, which in turn is well approximated by a Poisson distribution. As expected, larger deviations between the equivalent model's analytic solution and the approximation are noted for increasing values of $\hat{\lambda}_2$. In general, all the models in this range are well approximated by the equivalent model.

- The two cases presented in Figure 12 have $\hat{\lambda}_2 > 0.8$, for which the Gamma distribution is an approximation of the equivalent model's analytic solution. The distribution in some cases (see for instance the top figure), presents some deviations with respect to the equivalent model. However, globally a good agreement is obtained in most of the cases (failing ratio, based on a 0.5 maximum error is 21.7%).

- Two cases in an intermediate range $0.2 < \hat{\lambda}_2 < 0.8$ are shown in Figure 13. Again, the equivalent model's analytic solution is well representative of the distribution (failing ratio, based on a 0.5 maximum error is 3.2%). It is noted that for such values of $\hat{\lambda}_2$ an approximation of the equivalent model analytic solution is not available.

Thus, in most of the tested cases the equivalent model is able to catch the behaviour of a generic random GIA model, and thus represents a good approximation (global failing ratio, based on a 0.5 maximum error is 6%). In the cases where the equivalent model does not yield a good approximation, the internal structure of the $\mathcal{R}$ and $\mathcal{C}$ compartments become relevant and subsequent events that affect $n_R$ and $n_C$ become dependent on each other, and thus are non-Markovian.

**C. GIA Model for Large $\hat{\lambda}_R$.** To test the behaviour of a generic GIA model in case of large $\hat{\lambda}_R$, the GIA random models (generated as described in 2.B and analyzed in the main text) were modified by changing the process rates associated to the renewing compartment to achieve $\hat{\lambda}_R = 30$. To this aim, considering that infinite solutions are possible, we applied a global search method, and more specifically a Genetic Algorithm (10). We therefore setup an optimization problem, where the process parameters are the optimization variables and the cost function is the error of the current $\hat{\lambda}_R$ with respect to the target.

The envelope of curves obtained in all the random models and some illustrative profiles are shown in Figure 14. A reference Normal distribution, characterized by unitary mean and variance equal to $1/\hat{\lambda}_R = 1/30$ is also reported: this curve corresponds to the distribution expected in the equivalent model for which $\hat{\lambda}_1 = \hat{\lambda}_R$. Deviations become relevant, when the internal structure of compartments in a random model leads to subsequent events that are not independent from each other. These effects alter the variance of the Normal distribution. In fact, Figure 4 in the main text is based on the same simulation results, but in this case the rescaling is done considering both the mean number of cells and its variance (a Normal distribution is a two-parameter distribution).

**D. GIA$^B$ Test Case: bimodal distribution.** In the previous subsection we increased $\lambda_R$ in a way which assures that other parameters within $\mathcal{R}$ stay of the same order of magnitude. Here, we address the question what happens if some parameters within $\mathcal{R}$ are much smaller than parameters of $\mathcal{C}$, such as $\gamma_C$. For that purpose, we study another simple GIA model, let us call it GIA$^B$, as a modification of the GIA$^0$ test model defined by 14. In the GIA$^B$ model the renewing compartment is composed by two states $X_1$ and $X_2$, instead of only one. Cells in these states divide asymmetrically (i.e. one daughter cell remains within the renewing compartment while the other enters the committed compartment) or change state between $X_1$ and $X_2$ (*cell state switching*) while still remaining within the renewing compartment. The committed compartment of the system is composed just by a single state, $X_3$, and cells in this state either duplicate or die (as the previous state $X_2$ in Eq. 14). This corresponds to the model

$$X_1 \xrightarrow{\lambda_1} X_1 + X_3, X_2 \xrightarrow{\lambda_2} X_2 + X_3, X_1 \xrightarrow{\omega_{12}} X_2, X_2 \xrightarrow{\omega_{21}} X_1, X_3 \xrightarrow{\lambda_3} X_3 + X_3, X_3 \xrightarrow{\gamma} \emptyset. \qquad [44]$$

In this model, the effective parameters as defined in section 4.B, $\lambda_R = \lambda_1 P_1^* + \lambda_2 P_2^*$, where $P_i^* = \frac{\omega_{ji}}{\omega_{ij}+\omega_{ji}}$, $i, j = 1, 2, i \neq j$, and $\gamma_C = \gamma$. As before, we define the non-dimensionalized parameters $\hat{\lambda}_R = \lambda_R/\gamma_C$ and here we also define $\hat{\omega} = \omega_{12}/\gamma_C$, and further the parameter ratios $a = \lambda_1/\lambda_2$ and $b = \omega_{12}/\omega_{21}$. In the following we test this model for different values of $a$ and $\hat{\omega}$ as reported in Table 2, while fixing $\hat{\lambda}_R = 30$, which is our main scaling parameter, as well as $\hat{\lambda}_C = 0$ and $b = 1$.

The rescaled distribution of the number of cells in the committed compartment $\mathcal{C}$ (i.e. in state $X_3$), $n_C$, obtained at the final simulation time $\tau$, is shown in Figure 15. A value of $\tau$ equal to $20/\alpha_{\min}$ (where $\alpha_{\min}$ is the minimum of all rate parameters) was chosen to assure that the steady state is reached. Considering first the test cases GIA$^B$#1 and GIA$^B$#2 according to Table 2, which are characterized by $a = 1$ (i.e. there is no difference in the division timescales for the two renewing states), they both lead to a Normal distribution, independently on the value assumed by $\hat{\omega}$. Test cases GIA$^B$#3 to GIA$^B$#7 instead are all characterized by $a = 10$, and different orders of magnitude for $\hat{\omega}$ are tested. The distribution in these cases is Normal until $\hat{\omega} \geq \hat{\lambda}_R/10$ (see cases GIA$^B$#3 to GIA$^B$#5); when $\hat{\omega}$ is significantly lower than $\hat{\lambda}_R$, then bimodality emerges (see cases GIA$^B$#6 and GIA$^B$#7). Looking at the extreme case, GIA$^B$#7, cells in each renewing state, if analyzed independently, would result in a Poisson distribution in the committed compartment with different mean values (low for the slow-dividing state and large for the fast-dividing one). Thus, globally the distribution is in line with a bimodal distribution computed as

$$P(n) = \beta \text{Poisson}(\hat{\lambda}_R^{(1)}) + (1 - \beta)\text{Poisson}(\hat{\lambda}_R^{(2)}), \qquad [45]$$

in which $\beta$ is the mixing parameter, computed as

$$\beta = \frac{\bar{n} - \bar{n}_2}{\bar{n}_1 - \bar{n}_2}, \qquad [46]$$

and the parameters $\hat{\lambda}_R^{(i)}$ and $\bar{n}_i$ for $i = 1, 2$ correspond to the parameter $\hat{\lambda}_R$ and to the mean number of cells of a system in which the renewing compartment would be composed just by state $X_i$. The total mean number of cells is instead indicated by $\bar{n}$. The bimodal distribution given by Eq. 45 is indicated as a black dashed-dotted line in Figure 15.

### 5. Analysis of the Generalized Population Asymmetry Model

In the main text it is shown that GPA models predict asymptotically, for large times $t$, the same rescaled clone size distribution, that is, an Exponential distribution of unitary mean.

In Figure 16 the 50%tile distribution of all the GPA models analysed is shown at different levels of extinction (which are related to the different time points), showing a gradual convergence to the expected Exponential distribution.

Thus, the Markov approximation to all GPA models, Eq. 12 in the main text (the *equivalent model* of GPA models), becomes accurate for sufficiently large $t$ and no significant deviations are observed. This also means that for large $t$, the distribution is independent of the choice of parameters, since only the mean value of surviving clones, $\bar{n}_s$, depends on parameters, which however, does not affect the rescaled distribution in terms of $x = \frac{n}{\bar{n}_s}$. We can therefore abstain from an extended study of different parameter regimes. This is in contrast to the GIA model class where distributions depend sensitively in the choice of parameters if we are not in the scaling regime of large $\hat{\lambda}_R$, and the non-Markovian nature of GIA models can become relevant, as we showed in the previous section.

### 6. Asymptotic clone size distributions: Mathematical analysis

In the previous two sections we studied numerically how a Markovian representation can approximate general cell fate models (GIA and GPA) models. Here we study from an analytical view point how generic GIA and GPA models converge to the respective limiting distributions, for large time $t$ (GPA models) and large $\hat{\lambda}_R$ (GIA models).

Similar to section 4.B, we define $\boldsymbol{n}_R$ and $\boldsymbol{n}_C$ as the cell number vectors (here: actual cell numbers of the stochastic model, not mean cell numbers) restricted to the states of compartments $\mathcal{R}$ and $\mathcal{C}$, respectively. We further define the accumulated cell numbers $n_R = \sum_i (\boldsymbol{n}_R)_i$ and $n_C = \sum_i (\boldsymbol{n}_C)_i$ in $\mathcal{R}$ and $\mathcal{C}$, respectively. Considering $n_R$ and $n_C$ as *observables* of our compartment model, this corresponds to a *Hidden Markov Model* in that the dynamics of the observables are not Markovian, yet they are entirely determined by a set of states which follow a Markov process.

#### A. General dynamics of $C$-cells for GIA and GPA models.

**Comments on the effective rate parameter $\lambda_R$.** For general GIA and GPA models in the compartment representation of Eq. 9, main text, the effective rate parameter $\lambda_R$ (i.e. the frequency of cell divisions in $\mathcal{R}$ per cell), is defined similar as in section 4.B, yet, here we take into account that $\lambda_R$ can depend on time via the – not necessarily stationary – distribution of cells within $\mathcal{R}$ (since the process is non-Markovian). Hence, in these more general terms, we define $\lambda_R(t) = \sum_{i\in\mathcal{R}} \lambda_i P_i^R(t)$ where $P_i^R(t) = \frac{\bar{n}_i(t)}{\bar{n}_R(t)}$ is the probability of a single cell to be in state $i$ at time $t$. $P_i^R(t)$ may variate after each event $E$, as the conditional probability $P^R|_E$, provided that an event $E$ has just occurred, differs from the stationary state distribution.

In homeostasis, where the number of $R$-cells must on average stay constant, $\lambda_R$ is also the rate, per $R$-cell, at which $C$-cells are created from $R$-cells, via events $R \to R + C, R \to C + C$, or direct transition, $R \to C$. Thus, the total rate of $C$-cells being created from the $R$-cells by such events – let us call them $RC$-events – is $\lambda_R n_R$. While the non-Markovian nature of the process does not assure that such events are distributed exponentially, we can state that, by definition, the number of such creation events in a time period $\Delta t$, $N_{RC}$, has mean value $\langle N_{RC}(\Delta t)\rangle = \int_0^{\Delta t} \lambda_R(t) n_R(t)\, dt$.

While, $\lambda_R(t)$ may in principle depend on time, we note that when internal rates of $\mathcal{R}$ are fast compared to the time period $\Delta t$ above (an *internal rate* of $\mathcal{R}$ is a rate $\omega_{ij}$ where states $i, j$ are both in $\mathcal{R}$), then $\lambda_R(t)$ fluctuates quickly and we can make an adiabatic approximation, replacing $\lambda_R(t)$ by its average $\bar{\lambda}_R = \sum_{i\in\mathcal{R}} \lambda_i P_i^R$, where $P_i^{R*} = \frac{\bar{n}_i^*}{\bar{n}_R^*}$ is the steady state value of $P_i^R(t)$ (this corresponds for GIA models to the definition of $\lambda_R$ in section 4.B). This is fulfilled in our simulations of large $\hat{\lambda}_R$, since internal rates, such as $\hat{\omega}$ defined in section 4.D, scale with $\hat{\lambda}_R$ when $\lambda_R \to \infty$ (see section 4.C). Hence, the time scales of internal rates are substantially smaller than the relevant time scale $\Delta t = 1/\bar{\gamma}_C$, the lifetime of generated $C$-cells. Therefore, when comparing with simulation results, it is generally appropriate to assume that $\lambda_R(t) \approx \bar{\lambda}_R$ is constant. In the following subsection, we will discuss this case. The case when internal rates are slower than the time scale $\gamma_C$ is discussed in the subsequent subsection.

**Asymptotic distributions of $C$-cells** Each $C$-cell created by an $RC$-event initiates a sub-clone within $\mathcal{C}$, defined through its progeny, which then follows the dynamics of $\mathcal{C}$. Such sub-clones evolve independently of each other (a defining characteristic of branching processes (5)). Let us call the number of cells of a sub-clone created by an $RC$-event at time $t_i$, which evolves over time $t$, as $\xi_i(t)$[‡]. Therefore, the total number of cells in $\mathcal{C}$ is the sum of independent random numbers $\xi_i$,

$$n_C(t) = \sum_{i=1}^{N_{RC}} \xi_i(t) \qquad [47]$$

Note that the random numbers $\xi_i(t)$ are not identically distributed, since their statistics depend on the time point of the $i$-th $RC$-event. In particular, the mean value, $\bar{\xi}_i(t - t_i) = \langle \xi_i(t)\rangle$ and variance $\sigma_\xi^2(t - t_i) = \langle(\xi_i(t) - \bar{\xi}_i)^2\rangle$ depend on the time passed since the respective $RC$-event at time $t_i$. Thus, we cannot apply the central limit theorem in its original form to the sum of random numbers, Eq. 47. However, a variation of the central limit theorem states that sums of non-identically distributed random variables, $\sum_i \xi_i$, converge to normally distributed random variables, if mean and variance of $\xi_i$ are finite, *and* they fulfill *Lindeberg's condition* (11).

The (strict) Lindeberg's condition is said to be fulfilled for a sequence of random numbers $\xi_i$, $i = 1, ..., N$, if

$$\max_i \frac{\sigma_i^2}{\sigma_N^2} \to 0, \text{ for } N \to \infty \qquad [48]$$

where $\sigma_i^2 = \langle(\xi_i - \bar{\xi}_i)^2\rangle$ and $\sigma_N^2 = \sum_{i=1}^N \sigma_i^2$. If this is fulfilled, then $n_C = \sum_{i=1}^N \xi_i$ converges for $N \to \infty$ to a random variable that is normal distributed.

To show that the $\xi_i$ fulfill Lindeberg's condition, we note that $\xi_i(t - t_i)$ follow a sub-critical multi-type branching process, for which $\bar{\xi}_i(t) \to 0$ for $t \to \infty$, which is assured since the eigenvalues of the adjacency matrix of $\mathcal{C}$ are all negative (since dominant eigenvalues of all SCCs in $\mathcal{C}$ are negative (12)). For multi-type branching processes the variance $\sigma^2$ is proportional to the mean value, hence $\sigma_i^2(t - t_i) \sim \bar{\xi}(t - t_i)$. Therefore, $\sigma_i^2 \to 0$ for $t \to \infty$, hence it is bounded, i.e there exists $C > 0$ such that $\sigma_i^2(t) < C$ for all $t$. Furthermore, since initially, at $t = t_i$, $\bar{\xi}_i(t_i) = 1$, we know that there exist $t_1 > 0$ and $\delta > 0$ such that $\bar{\xi}_i(t) > \delta$ for $t - t_i < t_1$. Now we recall that, since here we assume the validity of the adiabatic approximation discussed in the

[‡] We denote two $RC$-events which happen at the same time via a symmetric division of type $R \to C + C$ by different indices $i$ and $i + 1$, yet with $t_i = t_{i+1}$

previous subsection, the number of $RC$-events within a time period $\Delta t$ is $N_{RC}(\Delta t) \sim \lambda_R \int_0^{\Delta t} n_R(t')\,dt'$. For generic $\lambda_R$, $N_{RC}$ is finite and thus is $\sigma_N$, since all $\sigma_i(t) \to 0$ for large $t$. However, for $\lambda_R \to \infty$ or $n_R \to \infty$, we get that $N_{RC}(t_1) \sim \bar{\lambda}_R n_R \to \infty$ and thus $\sigma_N^2 = \sum_{i=1}^{N_{RC}} \sigma_i^2(t) > N_{RC}\delta \to \infty$. On the other hand, all $\sigma_i^2 < C$, which means that all $\frac{\sigma_i^2}{\sigma_N^2} < \frac{C}{\sigma_N^2} \to 0$ for $\lambda_R \to \infty$ or $n_R \to \infty$. Hence, Lindeberg's condition is fulfilled if $\lambda_R \to \infty$ or $n_R \to \infty$ and thus, $n_C$ becomes normally distributed,

$$n_C(t) = \sum_i^{N_{RC}} \xi_i(t) \to \text{Normal}(\text{mean} = \bar{n}_C, \text{variance} \sim \bar{n}_C) \qquad [49]$$

The variance scales with $n_C$ since variances of independent random numbers add linearly and each $\sigma_i^2 \sim \bar{\xi}_i$. The exact value of $\bar{n}_C$ and the pre-factor of the variance of $n_C$ in this limit depend on the (non-Markovian) model details.

**Deviations from a Normal distribution in the asymptotic case** The arguments leading to Eq. 49 hold for large $\hat{\lambda}_R$ if the internal rates of $\mathcal{R}$ are comparable to $\bar{\lambda}_R = \sum_i \lambda_i \frac{\bar{n}_i^*}{\bar{n}_R^*}$, which is satisfied for all cases we sampled randomly for numerical simulations, see section 4.C. However, if internal rates of $\mathcal{R}$ are much smaller than $\lambda_R$, then the adiabatic approximation $P_i^R(t) \approx \frac{\bar{n}_i^*}{\bar{n}_R^*}$ does not apply and $\lambda_R(t)$ may vary slower than the time scale $1/\bar{\gamma}_C$. For example, consider a GIA model in which $\mathcal{R}$ can be decomposed into two sub-compartments, say $\mathcal{R}_1$ and $\mathcal{R}_2$, whereby any rates $\omega_{ij}, \omega_{ji}$ with $i \in \mathcal{R}_1, j \in \mathcal{R}_2$ have $\omega_{ij}, \omega_{ji} \ll \bar{\lambda}_R$, as the example discussed in section 4.D. Then, the single cell in $\mathcal{R}$ (note that always $n_R = 1$ in GIA models) may spend long time periods in $\mathcal{R}_1$ and $\mathcal{R}_2$ respectively. Now, if $\bar{\lambda}_{R_1} = \sum_{i\in\mathcal{R}_1} \lambda_i \frac{\bar{n}_i}{\bar{n}_{R_1}} \neq \sum_{i\in\mathcal{R}_2} \lambda_i \frac{\bar{n}_i}{\bar{n}_{R_2}} = \bar{\lambda}_{R_2}$, then, for time periods exceeding $1/\bar{\gamma}_C$, the effective asymmetric division rates are $\bar{\lambda}_{R_1}$ and $\bar{\lambda}_{R_2}$ respectively, and during these time periods the distribution of $n_C$ cells has mean $\bar{n}_C^{(1)} \sim \bar{\lambda}_{R_1}$ and $\bar{n}_C^{(2)} \sim \bar{\lambda}_{R_2}$ respectively. Hence, the total clone size distribution will be the mix of two Normal distributions with mean $\bar{n}_C^{(1)}$ and $\bar{n}_C^{(2)}$, respectively, i.e. a bimodal distribution. This scenario is discussed in section 4.D, for the specific case of two states in $\mathcal{R}$.

**B. GIA models.** In GIA models, the number of $R$-cells is conserved, and in particular, for clones, we have $n_R = 1$ for all times. Hence, the rate of $RC$-events is simply $\lambda_R$. Now, if internal rates are fast and $\lambda_R \to \infty$, then $n_C$ becomes normally distributed, as argued above. Hence, also $n = n_R + n_C = 1 + n_C$ follows a Normal distribution, with mean $n_C + 1$ instead.

Nonetheless, if internal rates are less than $\gamma_C$ then bimodal distributions may be observed, as discussed in section 4.D.

**C. GPA models.** The dynamics of GPA models read, in compartment formulation,

$$R \xrightarrow{\lambda_R} \begin{cases} R+R & \text{Pr. } r_{RR} \\ R+C & \text{Pr. } 1 - r_{RR} - r_{CC} \\ C+C & \text{Pr. } r_{CC} \end{cases}, \qquad [50]$$

$$R \xrightarrow{\omega_{RC}} C, \qquad C \xrightarrow{\lambda_C} C + C, \qquad C \xrightarrow{\gamma_C} \emptyset \qquad [51]$$

Since the dynamics of $R$-cells do not depend on $C$-cells, we can first consider the formers' dynamics separately. In homeostasis, where $\lambda_R r_{RR} = \lambda_R r_{CC} + \omega_{RC}$, we have thus for $R$-cells,

$$n_R \xrightarrow{\lambda_R r_{RR} n_R} n_R \pm 1 \qquad [52]$$

This is a simple continuous time branching process with two offspring; yet it is non-Markovian: subsequent events may be correlated, since each event imbalances the internal distribution $P_i^R$ of cells in the compartment $\mathcal{R}$. Yet, as for $C$-cells, we can write the number of $R$-cells as a sum of independent (but not identically distributed) random variables. Let us consider for each $R$-cells, born at time $t_i$, the random variable $\xi_i^R$ describing its "survival" state, i.e. $\xi_i^R = 1$ if that cell is still in $\mathcal{R}$, and $\xi_i^R = 0$ if that cell has left $\mathcal{R}$ via symmetric differentiation, $R \to C + C$ or direct transition, $R \to C$[§]. Since these events do not depend on other cells, the random numbers $\xi_i^R$ are independent of each other, and thus,

$$n_R(t) = \sum_{i=1}^{N_b(t)} \xi_i^R(t) \quad , \qquad [53]$$

is a sum of independent, not identically distributed random variables. Here, $N_b(t)$ is the total number of birth events occurring at rate $\lambda_R r_{RR} n_R$, $R \to R + R$, up to time $t$. Since $\xi_i^R(t) \le 1$ and $\xi_i^R(t = t_i) = 1$, we can argue analogue to above for Eq. 49 that the sequence of $\xi_i^R$ fulfills Lindeberg's condition and thus $n_R$ converges to a Normal distribution, whereby the mean value $\bar{n}_R = 1$ (since due to homeostasis the mean number is constant and the initial condition is $n_R(t = 0) = 1$). Hence, the probability to have $n_R$ cells in $\mathcal{R}$ is

$$P(n_R) \propto e^{-\frac{(n_R - 1)^2}{2\sigma_R^2}} \sim e^{-\frac{n_R^2}{2\sigma_R^2}} \text{ for } n_R \gg 1 \quad . \qquad [54]$$

[§]Essentially, the random numbers $\xi_i^R$ are the 'branches' of the branching process

However, here, the variance $\sigma_R^2$ is a random variable itself: Since the $\xi_i^R$ are independent, $\sigma_R^2 = \sum_{i=1}^{N_b(t)} \sigma_i^2$, where $\sigma_i^2 = \langle(\xi_i^R - \bar{\xi}^R)^2\rangle$, and where $N_b(t)$ is a random variable. The random numbers $\xi_i^R$ can only have the values $\xi_i = 1$ or $\xi_i^R = 0$ and they follow a simple death process, so for $\xi^R = 0$, it must be $\sigma_i^2 = 0$, while for $\xi_i^R = 1$, the variance must be finite, let's say, $\sigma_i^2 = \beta(t) > 0$ where $\beta$ can in principle depend on time, yet is not known (it depends on the non-Markovian details of the model). Hence,

$$\sigma_R^2 = \sum_{i=1}^{N_b(t)} \beta(t)\xi_i^R = \beta(t) n_R \qquad [55]$$

since the number of summands with $\xi_i^R = 1$ is the number of surviving $R$-cells, i.e. $n_R$. Substituting $\sigma_R^2 = \beta(t)n_R$ into Eq. 54 gives,

$$P(n_R) \sim e^{-\frac{n_R^2}{2\beta(t)n_R}} = e^{-\frac{n_R}{2\beta(t)}} \qquad [56]$$

This is an Exponential distribution with mean value $\bar{n}_R = \langle n_R\rangle = 2\beta(t)$. Finally, when we enforce normalisation of the probability distribution, we get,

$$P(n_R) = \frac{1}{\bar{n}_R(t)} e^{-\frac{n_R}{\bar{n}_R(t)}} \text{ for } n_R \gg 1 \ . \qquad [57]$$

Eventually, we also have to "add" the $C$-cells. Since for $t \gg 1$, also $n_R \gg 1$, individual events $n_R \to n_R \pm 1$ do not significantly affect the distribution of $R$-cells, $P_i^R = \frac{\bar{n}_i}{\bar{n}_R}$ (in contrast to the case of $n_R = 1$ for GIA models), and hence we can assume the adiabatic approximation discussed above, where $P_i^R \approx P_i^{R*}$ and thus $\lambda_R \approx const.$. Therefore, $C$-cells are distributed according to a Normal distribution with mean $\bar{n}_C$ and variance $\sigma_{n_2}^2 \sim \bar{n}_C \sim \lambda_R n_R$. As argued in the main text, the mean value of $n_R$, conditionend on survival of a clone, $n_R > 0$, must grow over time, without bound if $t \to \infty$. Therefore, we can generally assume that $n_R \gg 1$, and hence both $\bar{n}_C \sim n_R \to \infty$ and $\sigma_C^2 \sim n_R \to \infty$. However, if we express the clone size in form of a rescaled variable $x = \frac{n}{\bar{n}_s}$ ($\bar{n}_s$ is the mean of surviving clones) we can write $x = x_R + x_C$ with $x_R = \frac{n_R}{\bar{n}_s}$ and $x_C = \frac{n_C}{\bar{n}_s}$, and note that the rescaled standard width of the distribution of $x_C$, $\sigma_{x_C} = \frac{\sigma_C}{\bar{n}} \sim \frac{\sqrt{\bar{n}_C}}{\bar{n}_R + \bar{n}_C} \sim \frac{\sqrt{n_R}}{n_R}$ vanishes for $t \to \infty$. Therefore, $x_C$ is effectively a constant in that limit, $x_C \approx \bar{x}_C \propto x_R$. Hence, also $x = x_R + x_C \propto x_R$ and thus, the rescaled clone size, $x = \frac{n}{\bar{n}_s}$, is distributed according to an Exponential distribution (here: a probability density function) with unit mean, and after renormalisation, we get that

$$P(x) = e^{-x} \text{ for } t \to \infty. \qquad [58]$$

This distribution is indeed observed in all our simulations of GPA models for large $t$.


1. P Greulich, BD MacArthur, C Parigini, RJ Sánchez García, Stability and steady state of complex cooperative systems: a diakoptic approach. *Royal Soc. Open Sci.* **6**, 191090 (2019).
2. MW Hirsch, H Smith, Monotone dynamical systems. *Handb. differential equations: Ordinary Differ. Equations*, 57 (2006).
3. TH Cormen, *Introduction to Algorithms.* (MIT Press), (2009).
4. KJ Arrow, A "dynamic" proof of the Frobenuis-Perron theorem for Metzler matrices. *Probab. Stat. Math.*, 17–26 (1989).
5. P Haccou, P Jagers, VA Vatutin, *Branching Processes: Variation, Growth, and Extinction of Populations.* (Cambridge University Press, Cambridge), (2005).
6. J Bang-Jensen, GZ Gutin, *Digraphs: Theory, Algorithms and Applications.* No. August, pp. 59–60 (2007).
7. DT Gillespie, Exact stochastic simulation of coupled chemical reactions. *The J. Phys. Chem.* **81**, 2340–2361 (1977).
8. N Van Kampen, *Stochastic Processes in Physics and Chemistry.* Vol. 2, p. 464 (1981).
9. T Antal, PL Krapivsky, Exact solution of a two-type branching process: Clone size distribution in cell division kinetics. *J. Stat. Mech. Theory Exp.* **2010** (2010).
10. DE Goldberg, *Genetic Algorithms in Search, Optimization and Machine Learning.* (Addison-Wesley Longman Publishing Co., Inc., Boston, MA, USA), 1st edition, (1989).
11. P Billingsley, *Probability and Measure.* (John Wiley and Sons), p. Theorem 27.4 (1995).
12. KJ Åström, RM Murray, *Feedback Systems: An Introduction for Scientists and Engineers.* (Princeton Unversity Press), pp. 426–426 (2008).

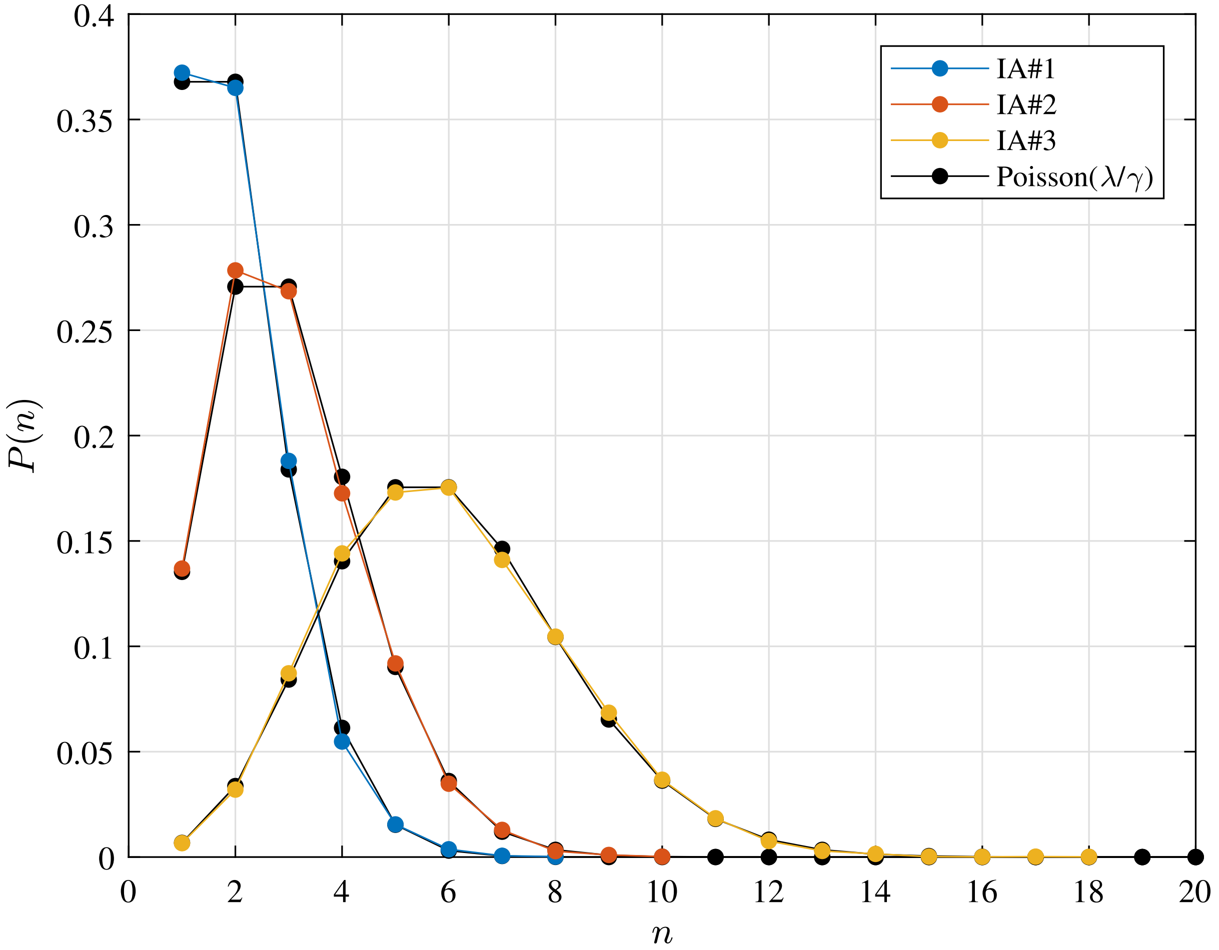


**Fig. 1.** Invariant Asymmetry (IA) test cases clone size distribution $P(n)$, that is the distribution of the total number of cells $n$ forming the progeny of a single initial cell in $\mathcal{R}$. For each case, the distribution is shown at $\tau$, which is well representative of the steady state condition. Tested parameters for cases IA$\#1$-$3$ are provided in Table 1; the numerical simulation results are compared to the expected Poisson distribution. The detailed discussion is reported in section 3.A.

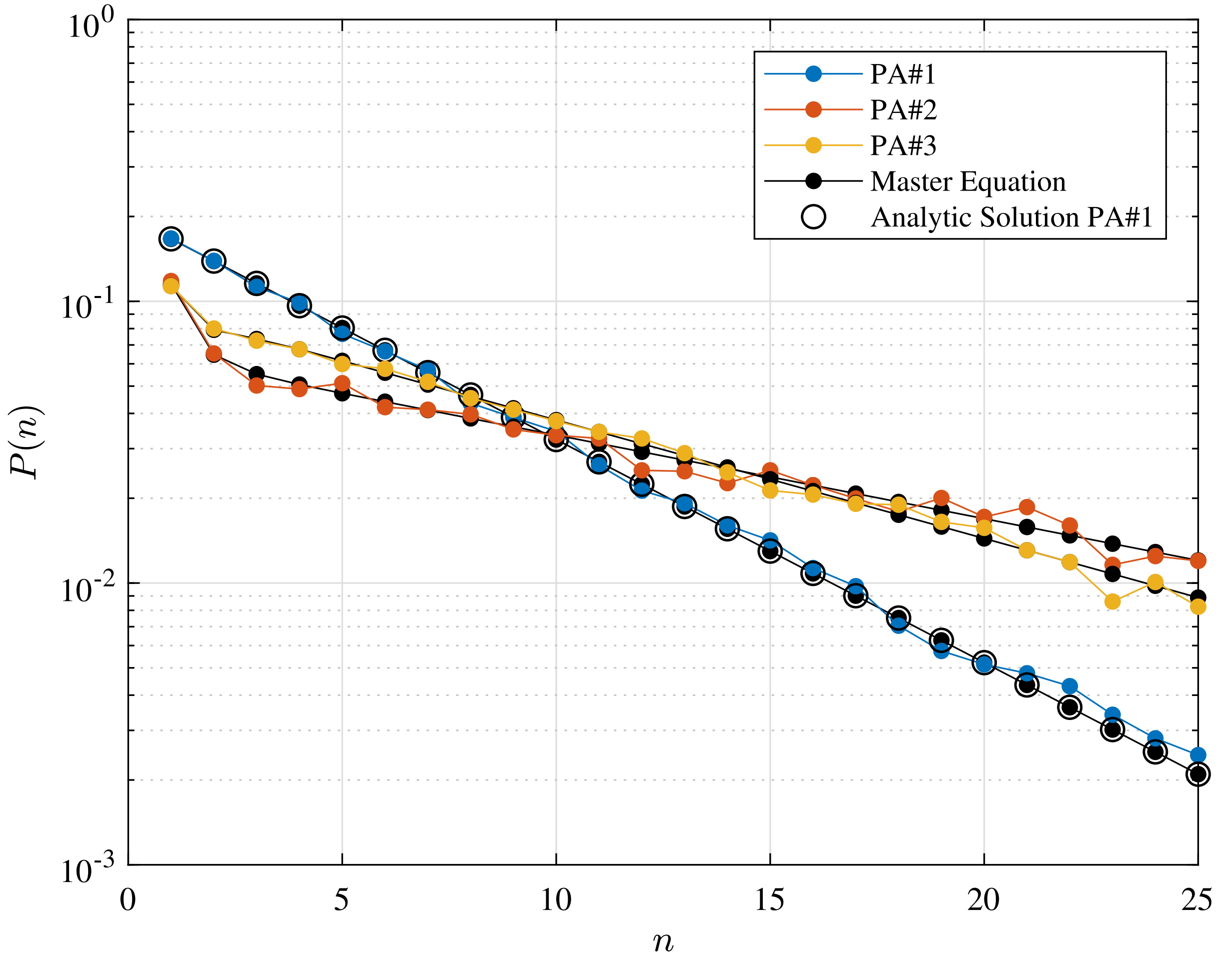


**Fig. 2.** Population Asymmetry (PA) test cases clone size distribution $P(n)$, that is the distribution of the total number of cells $n$ forming the progeny of a single initial stem cell. For each case, the distribution is shown at the final time $\tau$, at which the total extinction of the process is not yet achieved. Tested parameters for cases PA$\#1$-$3$ are provided in Table 1; the numerical simulation results are compared to the solution of the numerical integration of the master equation 8 and, for test case PA$\#1$, also to the reference analytic solution from Ref. (9). The detailed discussion is reported in section 3.A.

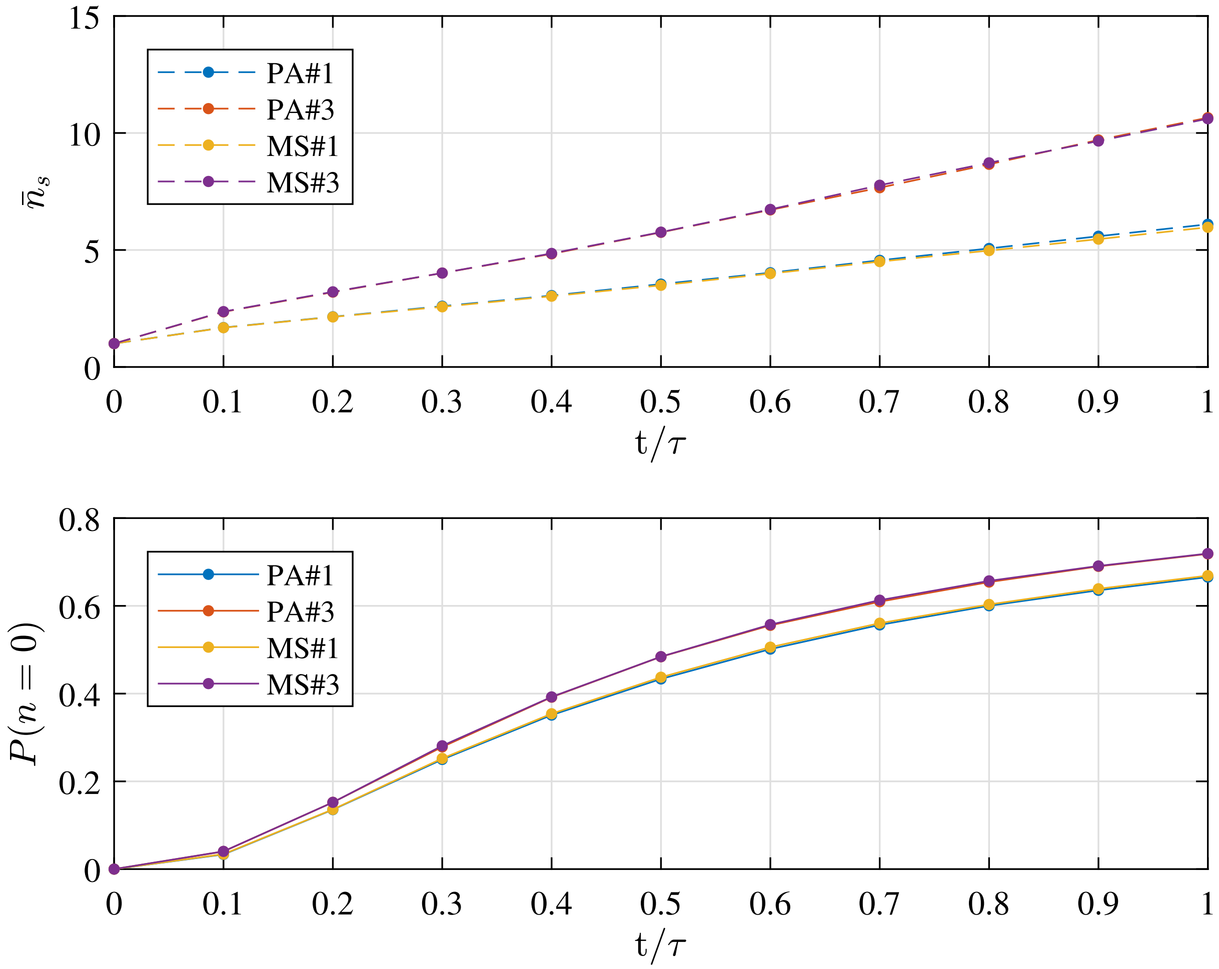


**Fig. 3.** Metastate (MS) test cases simulation results in terms of mean number of cells in the surviving clones $\bar{n}_s$ and extinction probability $P(n = 0)$ as function of time (scaled by the final simulation time $\tau$). As well as for the PA test cases, at $\tau$ the total extinction of the process is not yet achieved. Profiles from the numerical simulation for cases MS$\#1,3$ are compared to the corresponding PA$\#1,3$ test cases which are based on parameters provided in Table 1. The detailed discussion is reported in section 3.B.

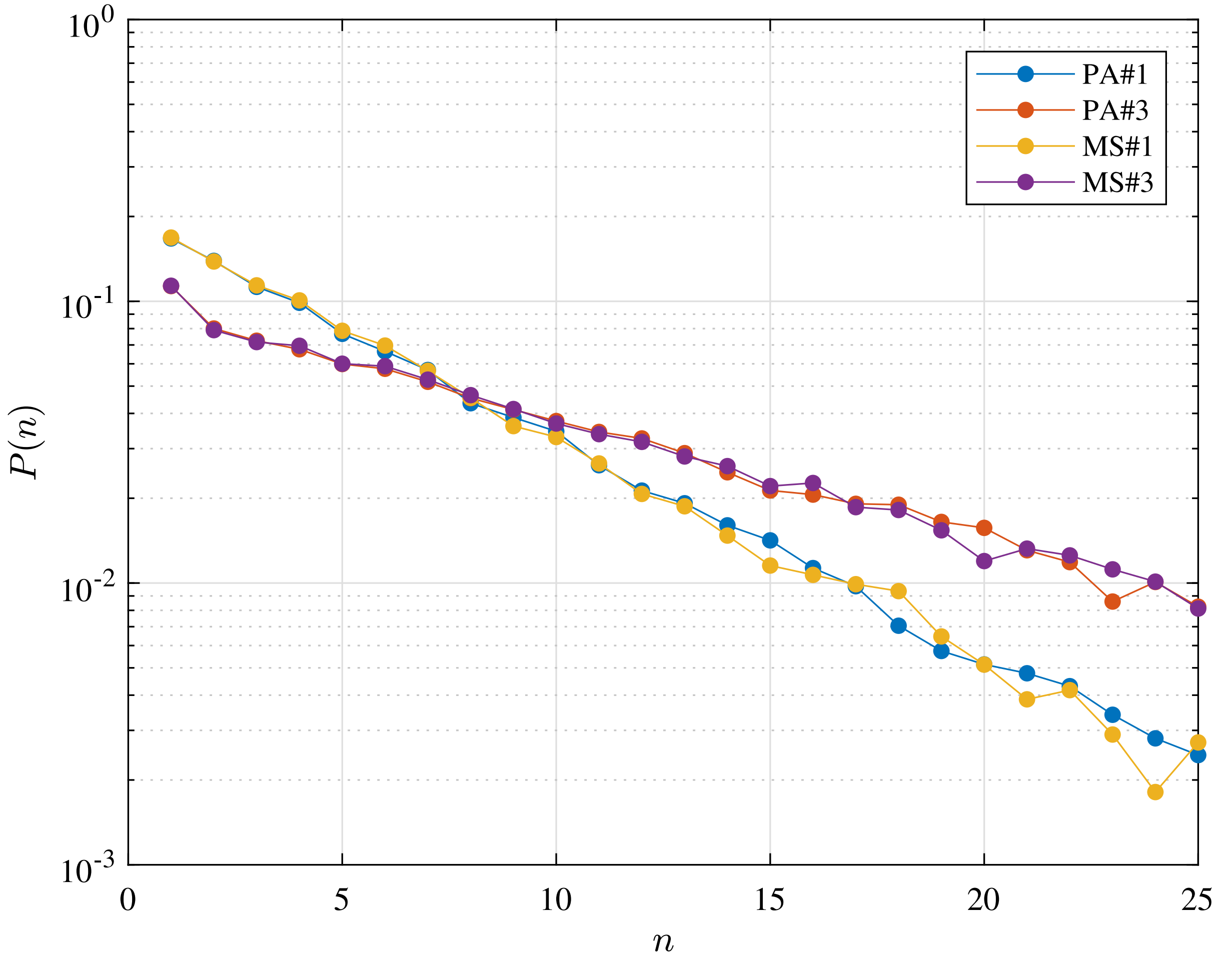


**Fig. 4.** Metastate (MS) test cases simulation results in terms clone size distribution $P(n)$, that is the distribution of the total number of cells $n$ forming the progeny of a single initial stem cell. As well as for the PA test cases, the distribution is shown at the final time, $\tau$, at which the total extinction of the process is not yet achieved. Profiles from the numerical simulation for cases MS$\#1,3$ are compared to the corresponding PA$\#1,3$ test cases which are based on parameters provided in Table 1. The detailed discussion is reported in section 3.B.

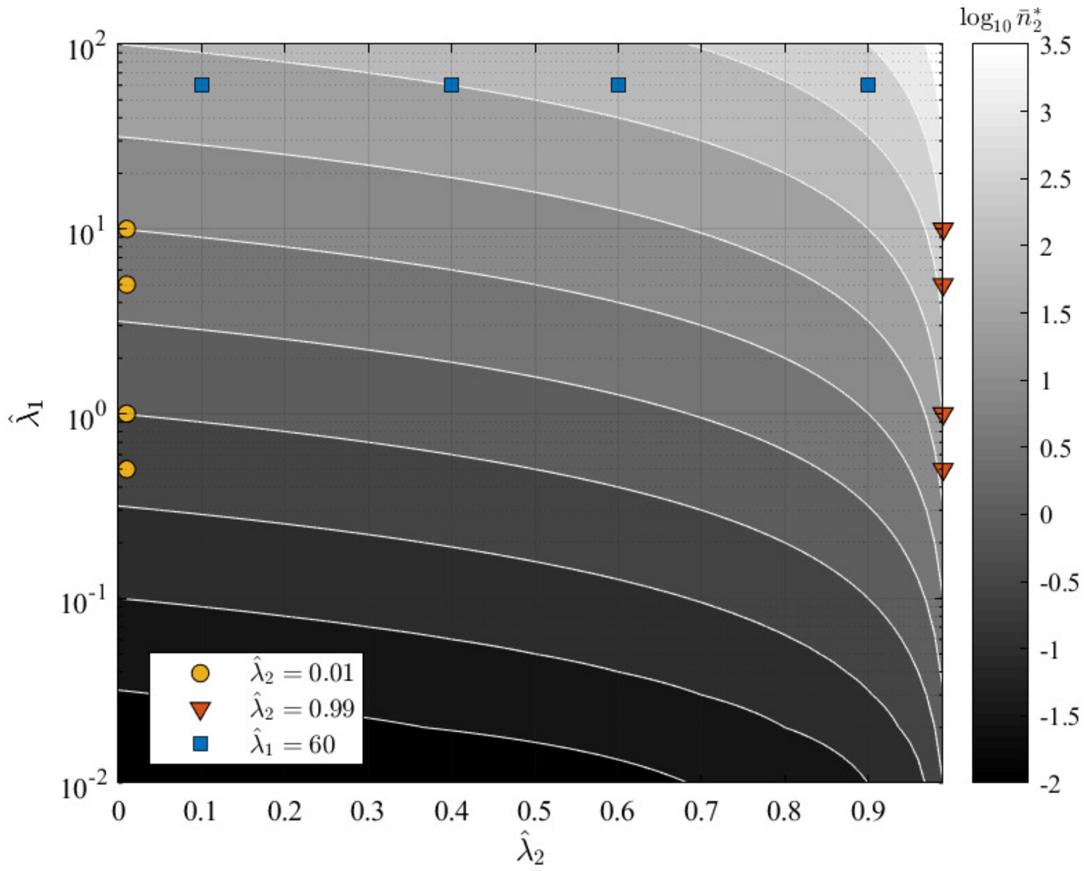


**Fig. 5.** GIA$^0$ test case parameters $\hat{\lambda}_1$ and $\hat{\lambda}_2$ over the contour map of the expected steady state mean number of cells in state $X_2$, $\bar{n}_2^*$. The tested conditions are divided in three groups representing the limiting behaviours discussed in in section 4.A, and for which the steady state distribution is shown respectively in Figure 6, Figure 7 and Figure 8.

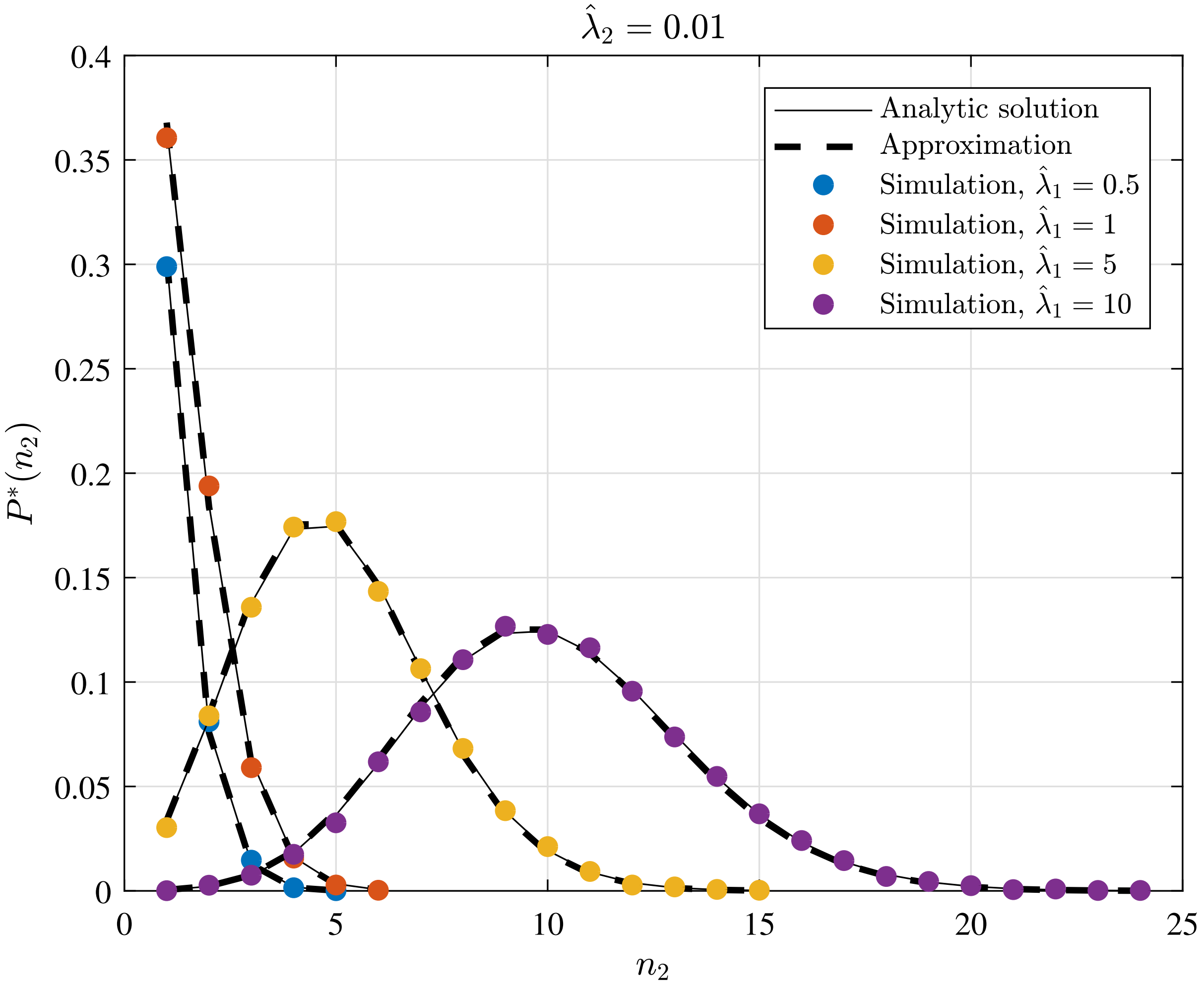


**Fig. 6.** GIA$^0$ test case (see section 4.A) results in terms of steady state distribution $P^*(n_2)$ of the the number of cells in state $X_2$, $n_2$. The tested parameters correspond to the condition $\hat{\lambda}_2 = 0.01$, as representative of the limiting case $\hat{\lambda}_2 \to 0$, and to different values of $\hat{\lambda}_1$. The results from the numerical simulations are compared to the analytic solution (Eq. 20), and its approximation, that is, the Poisson distribution (Eq. 27).

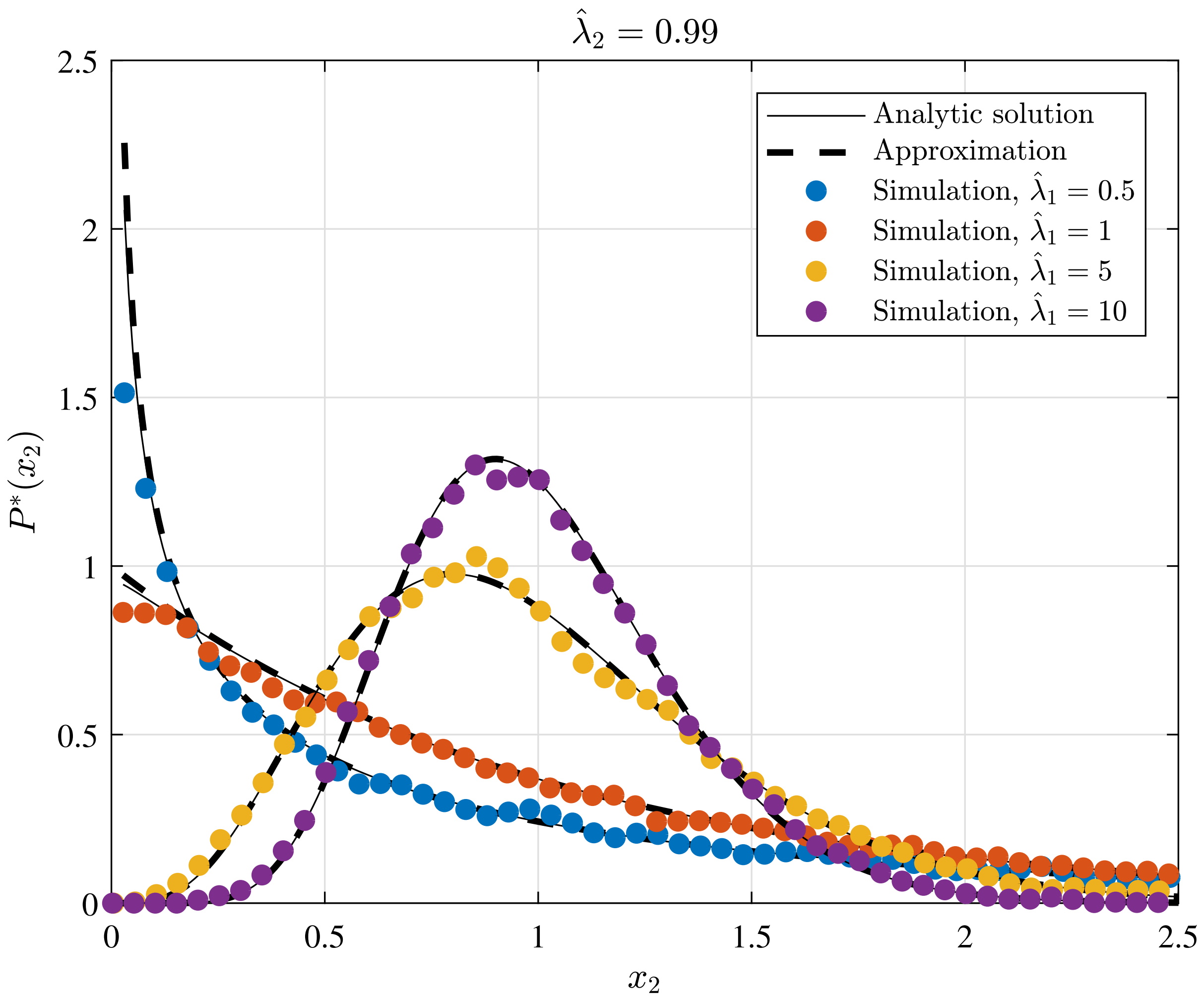


**Fig. 7.** GIA$^0$ test case (see section 4.A) results in terms of steady state rescaled distribution $P^*(x_2)$ of the the number of cells in state $X_2$, where $x_2 = n_2/\bar{n}_2^*$. The tested parameters correspond to the condition $\hat{\lambda}_2 = 0.99$, as representative of the limiting case $\hat{\lambda}_2 \to 1$, and to different values of $\hat{\lambda}_1$. The results from the numerical simulations are compared to the analytic solution (Eq. 23), and its approximation that is the Gamma distribution (Eq. 34).

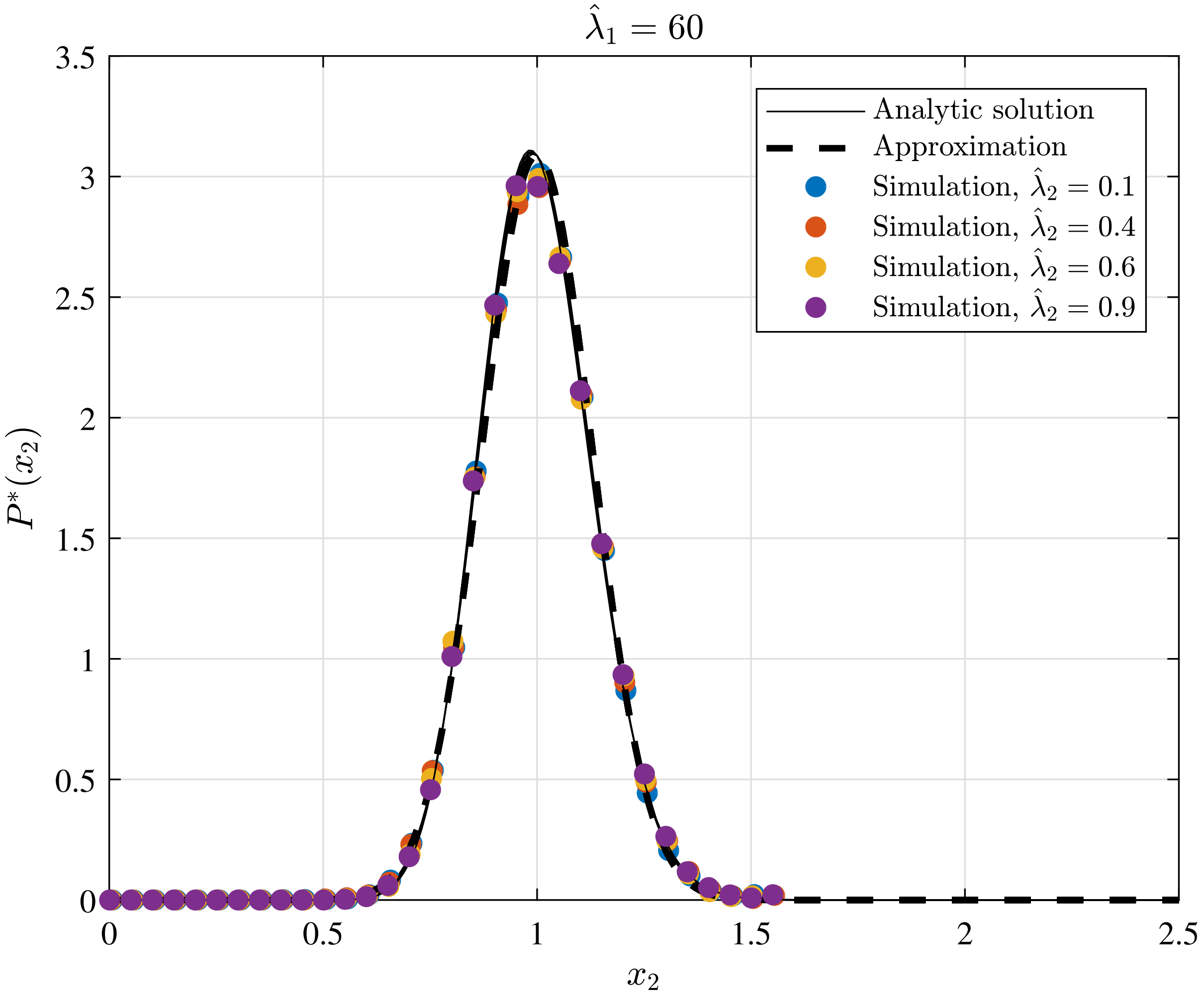


**Fig. 8.** GIA$^0$ test case (see section 4.A) results in terms of steady state rescaled distribution $P^*(x_2)$ of the the number of cells in state $X_2$, where $x_2 = n_2/\bar{n}_2^*$. The tested parameters correspond to the condition $\hat{\lambda}_1 = 60$, as representative of the limiting case $\hat{\lambda}_1 \to \infty$, and to different values of $\hat{\lambda}_2$. The results from the numerical simulations are compared to the analytic solution (Eq. 23), and its approximation that is the Normal distribution (Eq. 40).

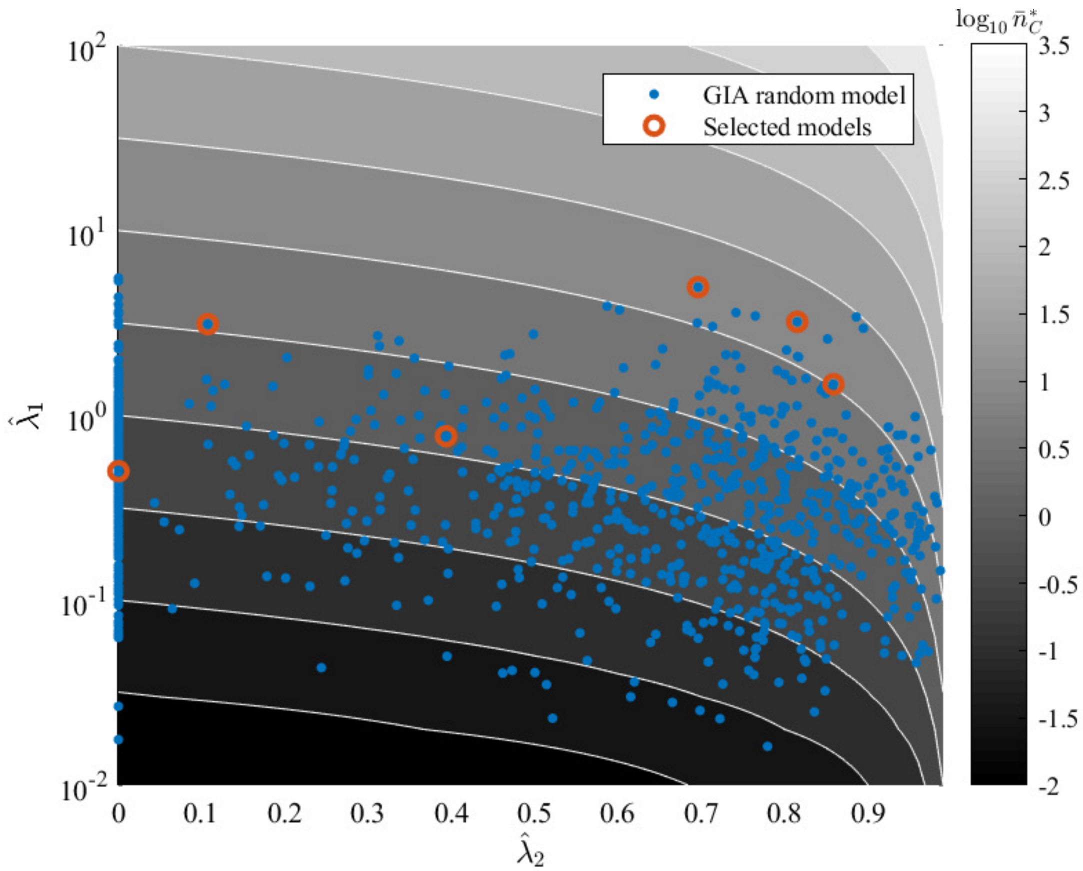


**Fig. 9.** GIA random models (generated as described in 2.B and analyzed in the main text) equivalent parameters $\hat{\lambda}_1 = \hat{\lambda}_R$ and $\hat{\lambda}_2 = \hat{\lambda}_C$ (see section 4.B) over the contour map of the expected steady state mean number of cells in the committed compartment, $\bar{n}_C^*$. Some illustrative cases, for which the steady state distribution is shown in Figure 11, Figure 12 and Figure 13, are highlighted.

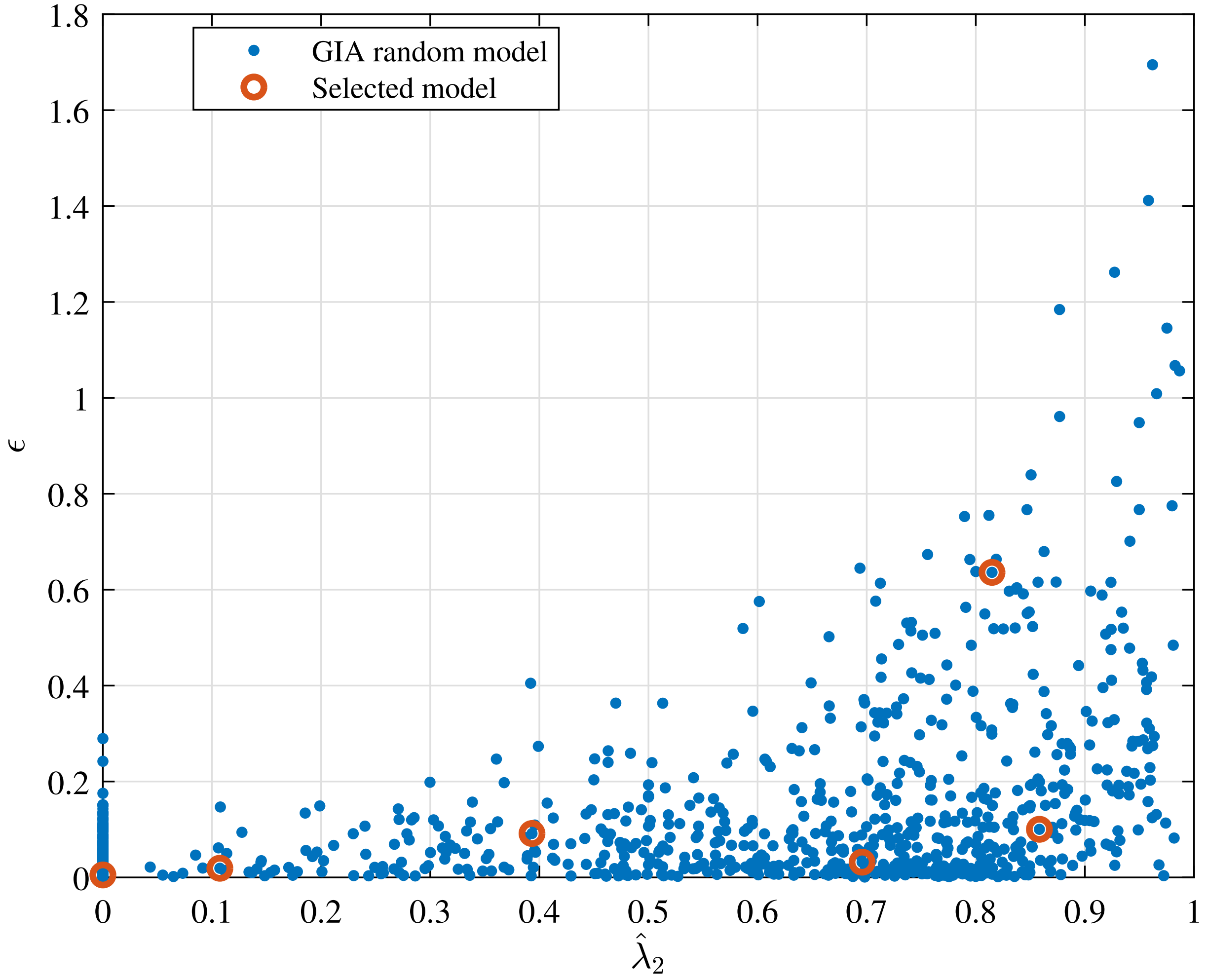


**Fig. 10.** Relative error of the the equivalent model approximation, $\epsilon$, (see definition in section 4.B) as function of $\hat{\lambda}_2 = \hat{\lambda}_C$ for the GIA random models (generated as described in 2.B and analyzed in the main text). The selected cases correspond to some illustrative cases for which the steady state distribution is shown in Figure 11, Figure 12 and Figure 13.

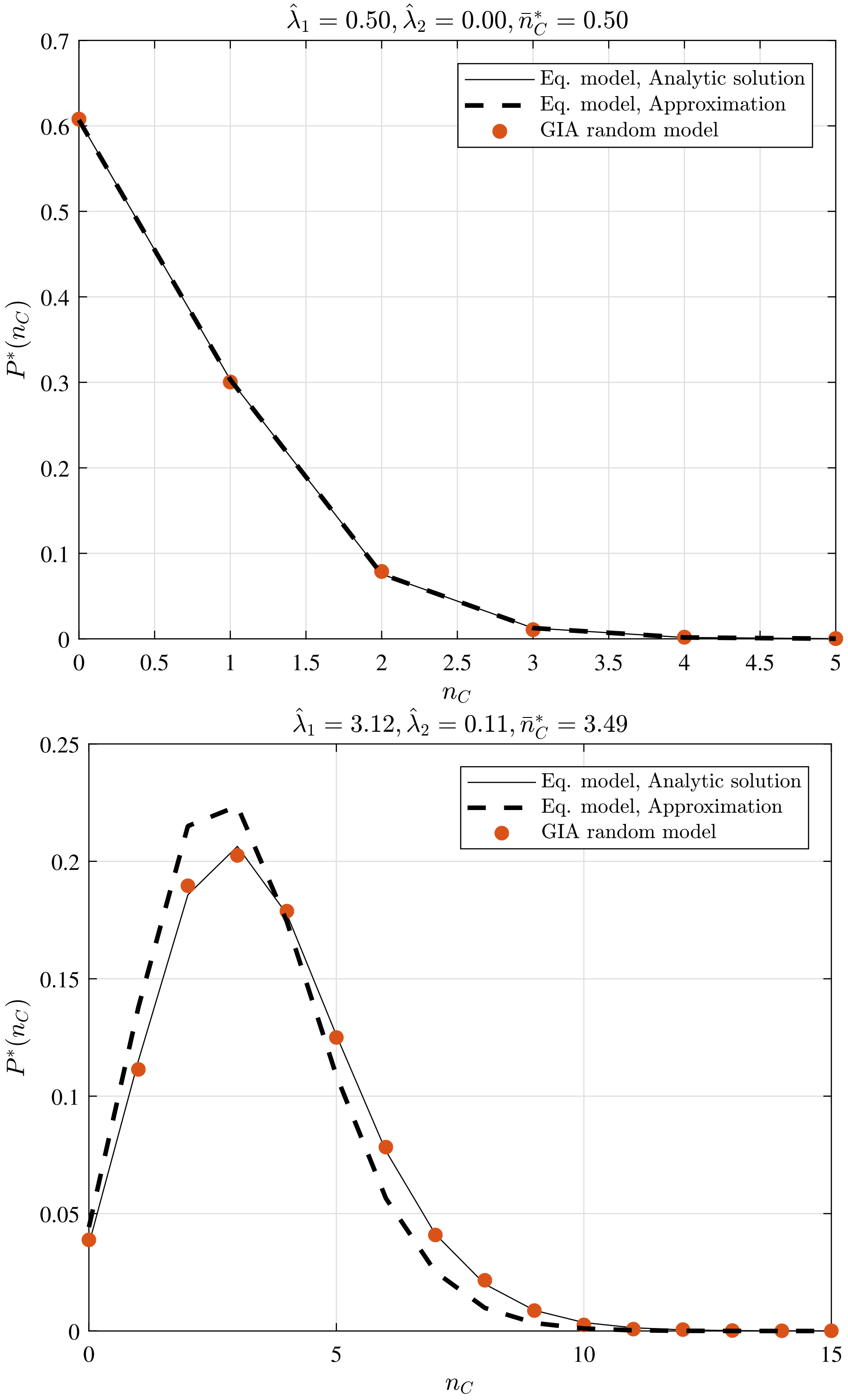


**Fig. 11.** GIA random models selected cases (see Figure 9 and 10) where $\hat{\lambda}_2 < 0.2$: the steady state distribution $P^*(n_C)$ of the number of cells in the committed compartment, $n_C$, is compared to that of the equivalent model (Eq. model in the legend) analytic solution and its approximation for low $\hat{\lambda}_2$ (i.e. the Poisson distribution, Poisson($\hat{\lambda}_1$)). Results discussion is reported in section 4.B.

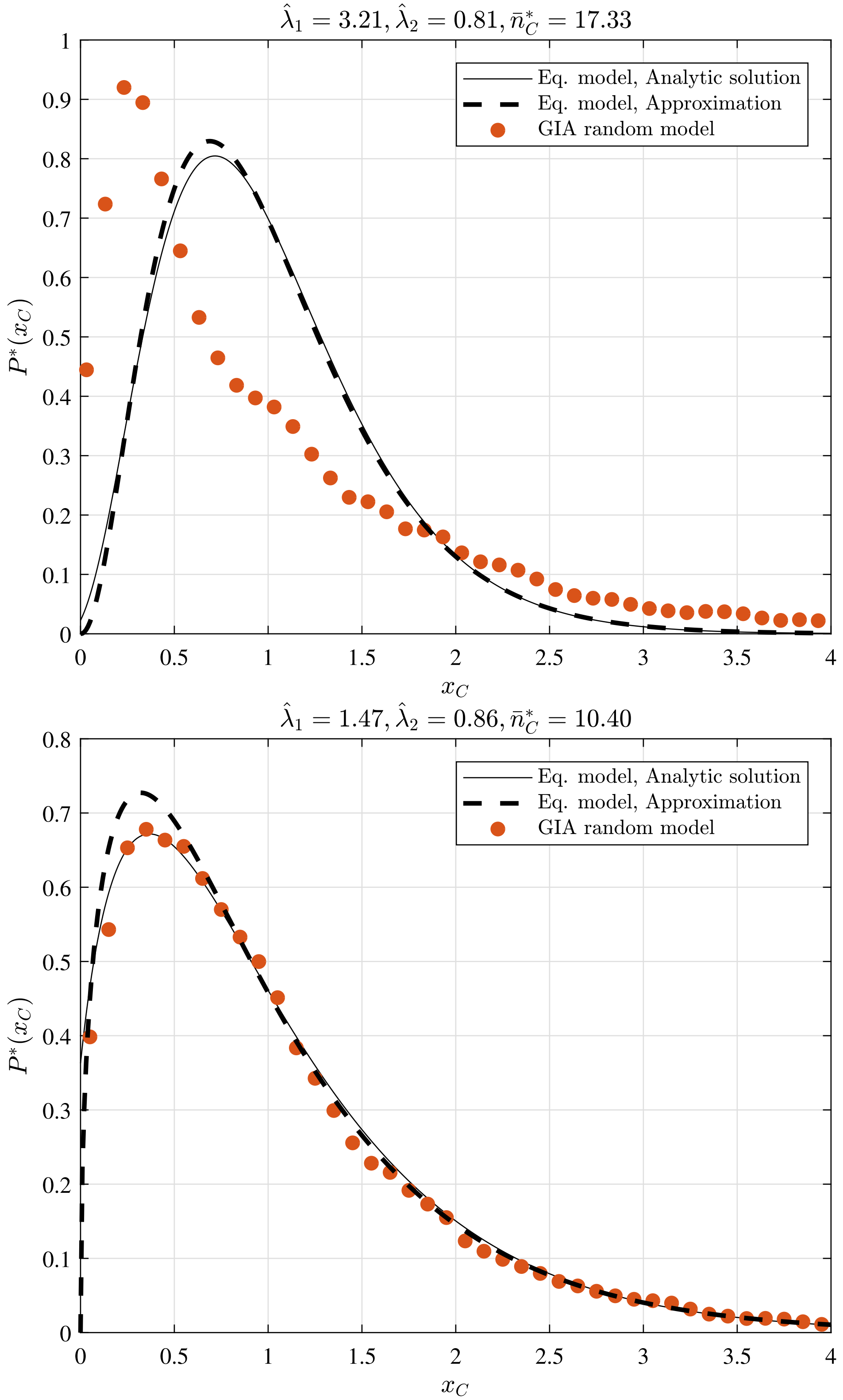


**Fig. 12.** GIA random models selected cases (see Figure 9 and 10) where $\hat{\lambda}_2 > 0.8$: the steady state rescaled distribution $P^*(x_C)$ of the number of cells in the committed compartment, where $x_C = n_C/\bar{n}_C^*$, is compared to that of the equivalent model (Eq. model in the legend) analytic solution and its approximation for high $\hat{\lambda}_2$ (i.e. the Gamma distribution, Gamma$(\hat{\lambda}_1, 1/\hat{\lambda}_1)$). Results discussion is reported in section 4.B.

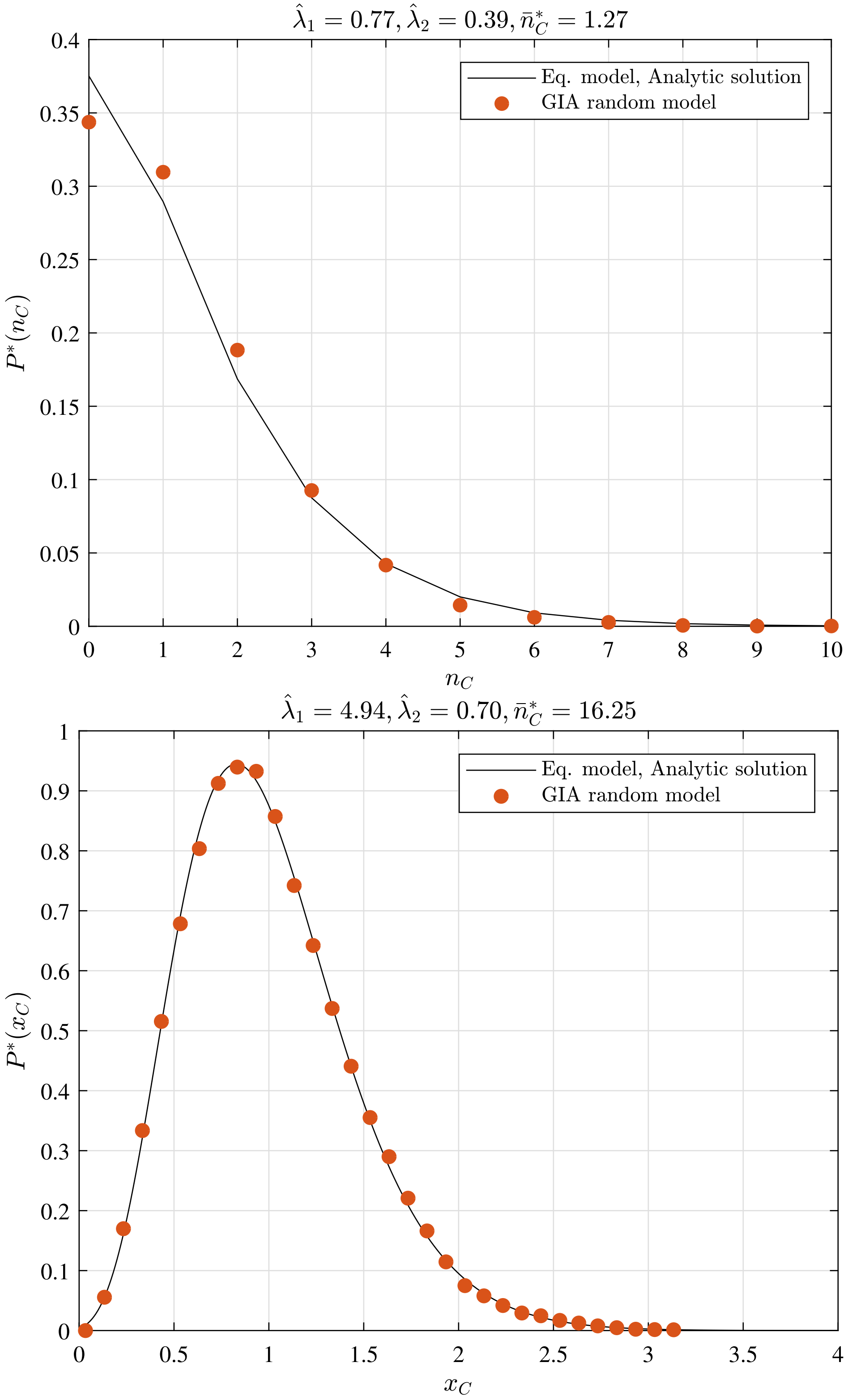


**Fig. 13.** GIA random models selected cases (see Figure 9 and 10) where $0.2 < \hat{\lambda}_2 < 0.8$: the steady state distribution $P^*(n_C)$ (or the rescaled distribution $P^*(x_C)$) of the number of cells in the committed compartment, $n_C$ (or in the rescaled case $x_C = n_C/\bar{n}_C^*$), is compared to that of the equivalent model (Eq. model in the legend) analytic solution. Results discussion is reported in section 4.B.

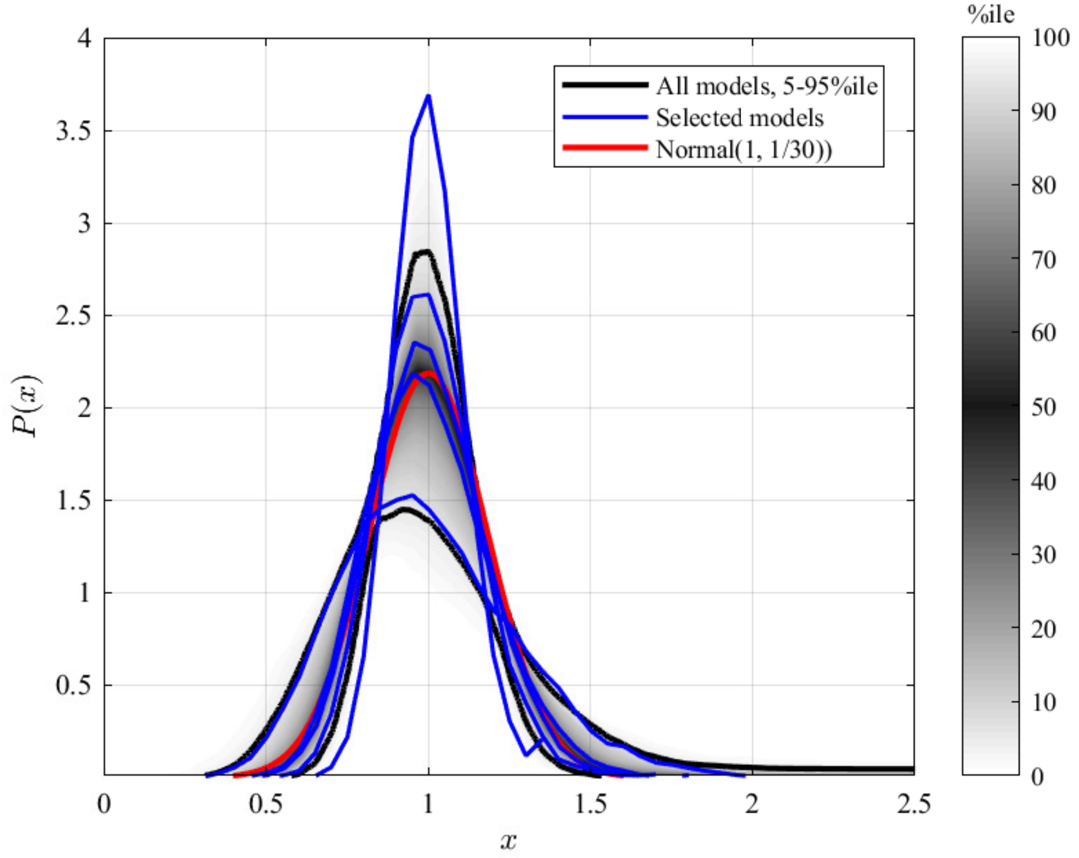


**Fig. 14.** Rescaled clone size distribution for the random GIA models when $\hat{\lambda}_R = 30$ at the final simulation time, which corresponds to $20/\alpha_{\min}$ ($\alpha_{\min}$ is the minimum process rate). The grey shade represents the percentile of all the simulations (black lines limit the 5-95%ile range); the blue curves correspond to some illustrative selected simulations. A reference curve corresponding to a Normal distribution of unitary mean and variance equal to $1/\hat{\lambda}_R = 1/30$ is shown in red. Distributions of the total number of cells $n$ are scaled by the mean number of cells $\bar{n}$, being $x = n/\bar{n}$. Simulations for which the final condition (20 times the inverse of the minimum process rate) is not achieved (due to computational limitations) are not included, resulting in 922 models. Results discussion in provided in section 4.C.

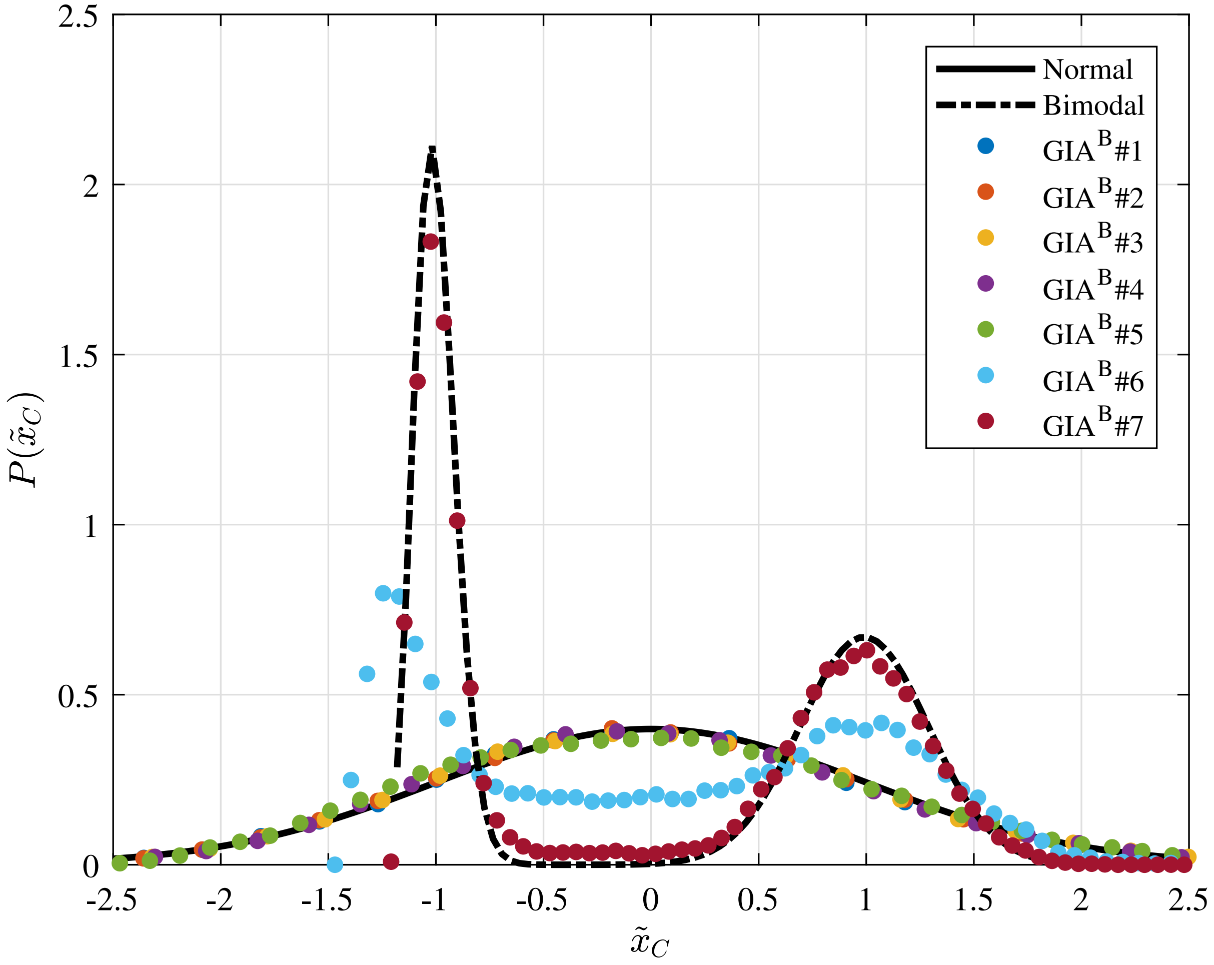


**Fig. 15.** Rescaled distribution of the cells number in the committed compartment in the $GIA^B$ test cases at time $\tau$, which is $20/\alpha_{\min}$ ($\alpha_{\min}$ is the minimum process rate). The distributions $P(\tilde{x}_C)$ of the number of cells in the committed compartment $n_C$ is rescaled considering that $\tilde{x}_c = (n_C - \bar{n}_C)/\sigma_{n_c}$, where $\sigma_{n_c}$ is the variance of $n_c$. In addition to the stochastic simulation results for different settings (see Table 2), the reference Normal and bimodal distributions are also shown. Results discussion is provided in section 4.D.

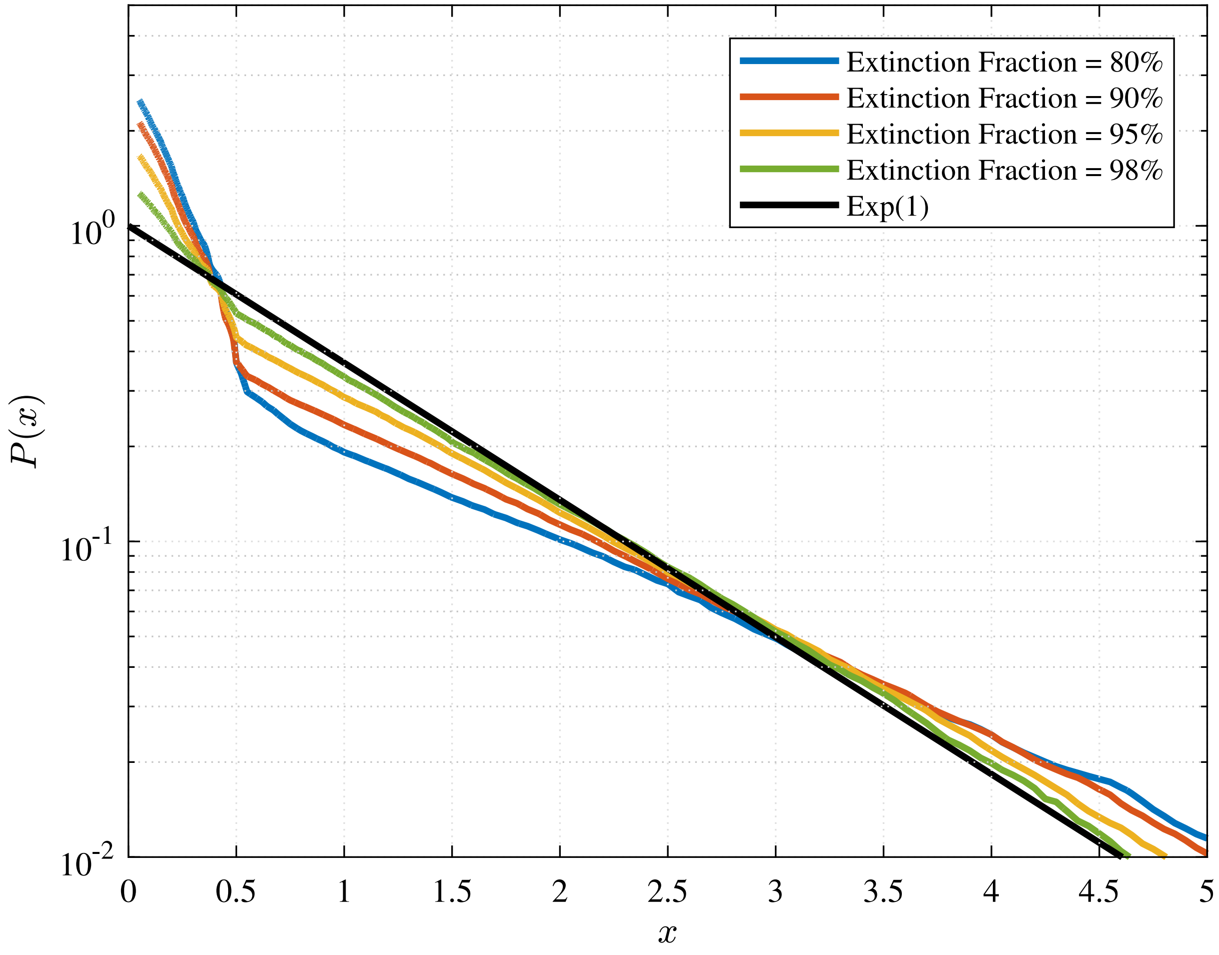


**Fig. 16.** Clonal size distribution (corresponding to the 50%ile curve) in the GPA random models at different extinction fraction (i.e. different time). The curves are compared to the expected Exponential distribution (see section 5).

**Table 1. IA and PA test cases simulation parameters (see section 3.A)**

| Case | $\lambda$ | $\gamma$ | r |
|---|---|---|---|
| IA#1 | 1.0 | 1.0 | - |
| IA#2 | 2.0 | 1.0 | - |
| IA#3 | 5.0 | 1.0 | - |
| PA#1 | 1.0 | 1.0 | 1/4 |
| PA#2 | 2.0 | 1.0 | 1/4 |
| PA#3 | 2.0 | 1.0 | 1/6 |

**Table 2. GIA$^B$ test case simulation parameters (see section 4.D)**

| Case | $\hat{\omega}$ | $\lambda_1/\lambda_2$ |
|---|---|---|
| GIA$^B$#1 | 3 $10^1$ | 1 |
| GIA$^B$#2 | 3 $10^{-2}$ | 1 |
| GIA$^B$#3 | 3 $10^2$ | 10 |
| GIA$^B$#4 | 3 $10^1$ | 10 |
| GIA$^B$#5 | 3 $10^0$ | 10 |
| GIA$^B$#6 | 3 $10^{-1}$ | 10 |
| GIA$^B$#7 | 3 $10^{-2}$ | 10 |
